# Atypical heat shock transcription factor HSF5 is critical for male meiotic prophase under non-stress conditions

**DOI:** 10.1101/2023.09.19.557986

**Authors:** Saori Yoshimura, Ryuki Shimada, Koji Kikuchi, Soichiro Kawagoe, Hironori Abe, Sakie Iisaka, Sayoko Fujimura, Kei-ichiro Yasunaga, Shingo Usuki, Naoki Tani, Takashi Ohba, Eiji Kondoh, Tomohide Saio, Kimi Araki, Kei-ichiro Ishiguro

## Abstract

Meiotic prophase progression is differently regulated in males and females. In males, pachytene transition during meiotic prophase is accompanied by robust alteration in gene expression. However, how gene expression is regulated differently to ensure meiotic prophase completion in males remains elusive. Herein, we identified HSF5 as a male germ cell-specific heat shock transcription factor (HSF) for meiotic prophase progression. Genetic analyses and single-cell RNA-sequencing demonstrated that HSF5 is essential for progression beyond the pachytene stage under non-stress conditions rather than heat stress. Chromatin binding analysis *in vivo* and DNA-binding assays *in vitro* suggested that HSF5 binds to promoters in a subset of genes associated with chromatin organization. HSF5 recognizes a DNA motif different from typical heat shock elements recognized by other canonical HSFs. This study suggests that HSF5 is an atypical HSF that enforces the gene expression program for pachytene transition during meiotic prophase in males.

## INTRODUCTION

Meiosis, which produces sperm and oocytes, is critical for germ cell development. Meiotic entry is followed by the meiotic prophase, in which meiosis-specific chromosomal events such as chromosome axis formation, homolog synapsis, and meiotic recombination occur sequentially (Handel and Schimenti 2010) (Zickler and Kleckner 2015) (Hunter 2015) (Page and Hawley 2004). The meiotic prophase is regulated by sexually dimorphic mechanisms, such that the gene expression program is altered for the subsequent developmental program of sperm production or oocyte maturation. In the male meiotic prophase, developmental progression beyond the pachytene stage is a critical event in which multiple gene regulatory programs inactivate gene expression on sex chromosomes, suppress transposable elements, and progress toward the post-meiotic stage (Li et al. 2013) (Sin et al. 2015) (da Cruz et al. 2016) (Ernst et al. 2019). Accordingly, a subset of male germ cell-specific transcription factors is required for progression beyond the pachytene stage (Bolcun-Filas et al. 2011) (Li et al. 2013) (Oji et al. 2020) (Horisawa-Takada et al. 2021) (Oura et al. 2021) (Xu et al. 2022) (Galan-Martinez et al. 2022) (Cecchini et al. 2023). However, it remains unclear how pachytene progression is ensured in the male meiotic prophase and which transcription factors are responsible for this process.

Previously, we identified MEIOSIN, which plays an essential role in meiotic initiation in males and females (Ishiguro et al., 2020). MEIOSIN, together with STRA8 (Kojima et al.2019), activates numerous meiotic genes. Among the target genes of MEIOSIN whose functions are unknown, we screened new germ cell-specific factors involved in meiosis (Takemoto et al. 2020) (Horisawa-Takada et al. 2021) (Tanno et al. 2022). In the present study, we identified the Heat shock transcription factor family 5 (*Hsf5*) gene that encodes a HSF as one of the MEIOSIN/STRA8 target genes.

The HSF family comprises several paralogs: HSF1, HSF2, HSF4, HSF5, and HSFY, which are common in humans and mice, while humans and mice additionally possess HSFX and HSF3, respectively (Akerfelt et al. 2010a). The mammalian HSF family drives gene regulation events that activate or repress transcription during stress responses and under non-stress conditions (Akerfelt et al. 2010a) (Abane and Mezger 2010) (Gomez-Pastor et al. 2018). Heat stress induces heat shock response (HSR), which is mediated by heat shock proteins (HSPs). The best known HSF paralog associated with the stress response is HSF1, which is present in the cytoplasm as a monomer when inactive and, upon sensing stress, forms a homotrimer that trans-locates into the nucleus, binds specifically to the heat shock element (HSE) in the genome, and regulates HSP gene transcription (Sarge et al. 1993) (Baler et al. 1993) (Wu 1995). Under non-stress conditions, the HSF family is known to regulate developmental processes other than stress responses (Abane and Mezger 2010). *Hsf1* knockout (KO) males produce fewer sperms compared to wild-type (WT) mice but are still fertile (Salmand et al. 2008). *Hsf2* KO males exhibit reduced spermatogenesis but are still fertile (Wang et al. 2003) (Kallio et al. 2002). *Hsf1* and *Hsf2* double knockout causes male infertility in mice (Wang et al. 2004). These suggested that HSF1 and HSF2 play synergistic roles in spermatogenesis under non-stress conditions.

Previous genetic studies have suggested that HSF5 homologs are involved in spermatogenesis in various species. In zebrafish, *Hsf5* mutant males were infertile with reduced sperm count, increased sperm head size, and abnormal tail architecture, whereas females remained fertile (Saju et al. 2018). In human testes, patients with azoospermia and low modified Johnson scores were associated with low expression of HSF5 (Chalmel et al. 2012). Additionally, although it has been demonstrated that disruption of *Hsf5* led to apoptosis during spermatogenesis in mice (Barutc et al. 2023), the precise mechanisms by which HSF5 is involved in meiosis remain elusive. Also, it remained elusive whether and how HSF5 plays overlapping and/or different roles to other HSFs in meiotic prophase progression in the testis. Furthermore, despite the presence of a DNA-binding domain, it is yet to be determined whether HSF5 acts as a transcription factor under stress or non-stress conditions like other HSFs.

Here, we show that mouse HSF5 plays an essential role in the meiotic prophase progression in male germ cells under non-stress conditions. Our single-cell (sc) RNA-sequencing (seq) analysis of *Hsf5* KO spermatocytes demonstrated that HSF5 was required for progression beyond the pachytene stage during spermatogenesis. Furthermore, chromatin immunoprecipitation sequencing (ChIP-seq) of HSF5 *in vivo* combined with DNA-binding analysis *in vitro* demonstrated that HSF5 binds to the promoters of a subset of genes whose biological functions are associated with chromatin organization through a DNA motif that is different from the typical HSE. The present study suggests that HSF5 acts as an atypical HSF under non-stress conditions that execute the gene expression program for pachytene transition during male meiotic prophase.

## RESULTS

### *Hsf5* is expressed during the meiotic prophase in the testis

Previously, we identified a germ cell-specific transcription factor, MEIOSIN, that directs the initiation of meiosis (Ishiguro et al. 2020). Our previous ChIP-seq analysis suggested that MEIOSIN and STRA8 bind to numerous genes in preleptotene spermatocytes whose functions are unknown. *Hsf5* was identified as one of the MEIOSIN/STRA8-bound genes (Fig. 1A). Our previous RNA-seq analysis suggested that the expression of *Hsf5* was one of the downregulated genes in *Meiosin* KO testes at postnatal day 10 (P10) when a cohort of spermatocytes underwent the first wave of meiotic entry (Ishiguro et al. 2020). However, the biological function of HSF5 has been unknown in mice.

**Figure 1.**
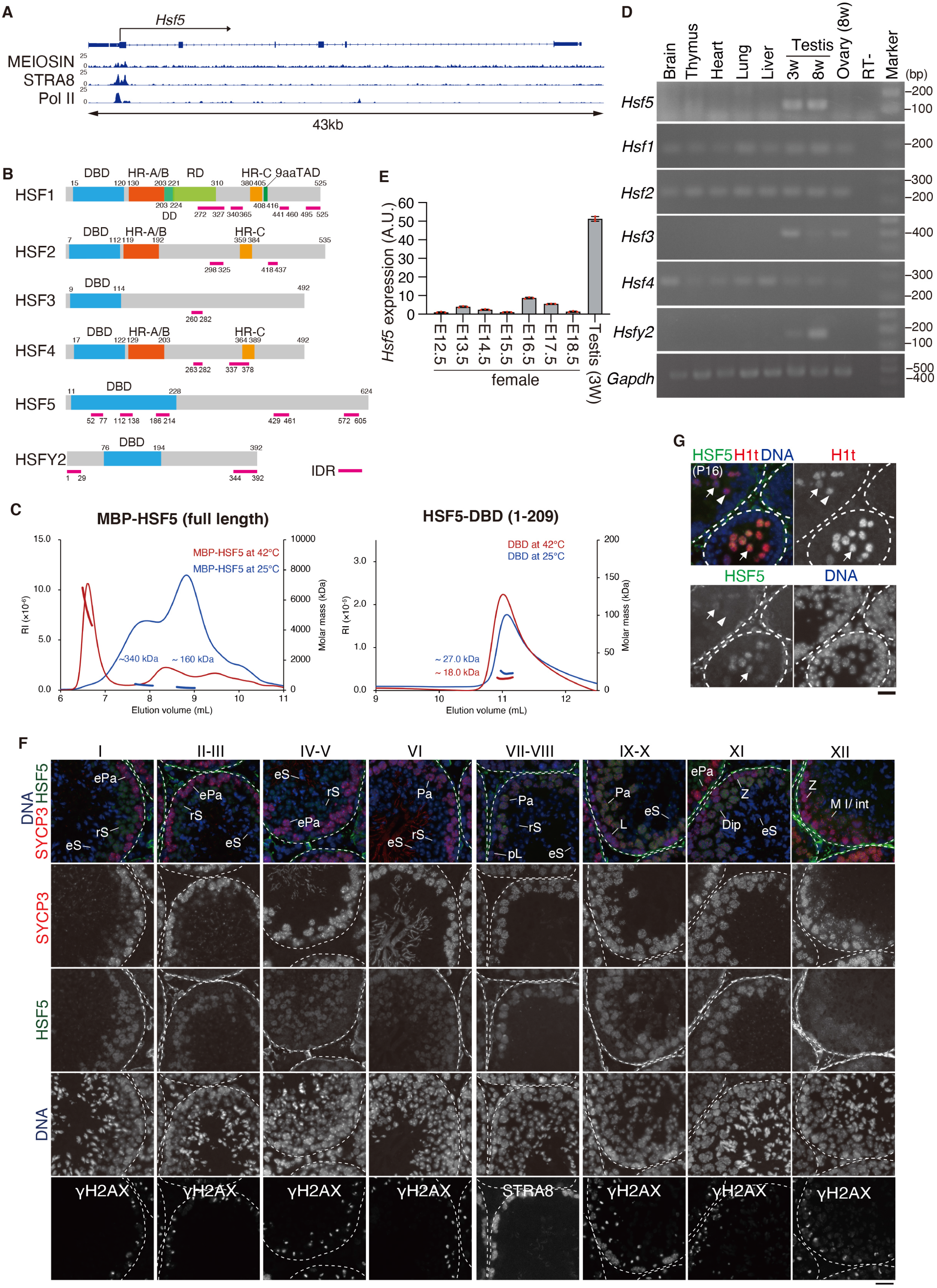
HSF5 is expressed during meiotic prophase in the mouse testis. **(A)**Genomic view of MEIOSIN and STRA8 binding peaks over the *Hsf5* locus. Genomic coordinates derived from NCBI. To specify testis specific transcription, RNA polymerase II ChIP-seq in the testis are shown (Li et al. 2013). **(B)** Schematic diagram of domain structure in six members of mouse heat shock factor (HSF) protein family. Domain name and the number of the amino acid residues are assigned according to Uniprot. DBD; DNA-binding domain. HR; heptad repeat. IDR; Intrinsically disordered region. DD; D domain. RD; Regulatory domain. 9aaTAD; Transactivation motif. **(C)** SEC-MALS profiles of MBP-full length HSF5 (left) and HSF5-DNA binding domain (a.a. 1-209) (right) at room temperature (blue) and after heat treatment at 42 °C for 30 min (red). Thin and bold lines show the refractive index (RI) profile and mass plot, respectively. **(D)** The tissue-specific expression pattern of *Hsf5* was examined by RT-PCR. Testis RNA was obtained from 3-weeks old (3w) and 8-weeks old (8w) male mice. Ovary RNA was obtained from adult 8-weeks old (8w) female mice. RT-indicates control PCR without reverse transcription. The data was acquired from two separate experiments. **(E)** The expression patterns of *Hsf5* in the embryonic ovary (E12.5-E18.5) and testis (3w) were examined by RT-qPCR. **(F)** Seminiferous tubule sections in WT testis (8-weeks old) were immunostained as indicated. pL: preleptotene, L: Leptotene, Z: Zygotene, ePa: early Pachytene, P: Pachytene, M I: Metaphase I, int: Interkinesis, rS: round Spermatid, eS: elongated Spermatid. Boundaries of the seminiferous tubules are indicated by white dashed lines. Roman numbers indicate the seminiferous tubule stages. Biologically independent mice (N=3) were examined in three separate experiments. Scale bar: 25 μm. **(G)** Seminiferous tubule sections in WT testis (P16) were immunostained as indicated. Arrow and arrowhead indicate HSF5-positive/H1t-positive and HSF5-negative/H1t-positive pachytene spermatocyte, respectively. Scale bar: 25 μm.

Six paralogous genes encoding HSFs were identified in the mouse genome (Fig.1B). The HSF family possesses a DNA-binding domain and acts as a transcription factor in the heat stress response and under non-stress conditions (Akerfelt et al. 2010a) (Abane and Mezger 2010). HSF5 possesses a winged-helix-turn-helix (WHTH)-like DNA-binding domain (Fig.1B). While HSF1, HSF2, and HSF4 possess two heptad repeats, HR-A and HR-B, that are predicted to form inter-molecular leucine zippers for homotrimer oligomerization (Rabindran et al. 1993), HSF5 lacks these heptad repeats (Fig. 1B). Size exclusion chromatography-multi angle light scattering (SEC-MALS) analysis indicates that the purified His-MBP-fused full length HSF5 protein was eluted at the peaks of approximate molar mass of 160 kDa and 340 kDa at 25 °C (Fig. 1C), suggesting that HSF5 exists as a mixture of monomer and dimer/trimer at 25 °C. SEC-MALS of HSF5 after 42 °C treatment resulted a peak with megadalton-scale size, suggesting that HSF5 forms high-order oligomers at higher temperature. This thermal response of HSF5 resembles HSF1 that forms higher-order oligomers upon heat shock (Kawagoe et al. 2022). In contrast, the HSF5 fragment (aa 1-209) containing DNA binding domain (HSF5-DBD) was in a monomeric state at both 25 °C and 42 °C, suggesting that oligomerization of HSF5 is mediated by other region than DBD. Thus, HSF5 exhibits a structural feature to undergo oligomerization *in vitro*.

We examined the expression patterns of *Hsf5* in different mouse tissues using RT-PCR. *Hsf5* mRNA was specifically expressed in the juvenile and adult mouse testes but not in other adult organs (Fig. 1D), which is in stark contrast to the ubiquitous expression profiles of *Hsf1, Hsf2,* and *Hsf4* (Fig. 1D, Fig. S1A). Similarly, *Hsf2* is expressed in the juvenile and adult testes. Although the expression profile of mouse *Hsf3* in any organ is unknown in the available database (Fig. S1A), our RT-PCR suggested *that Hsf3* is expressed in the testes and ovaries. The spermatogenic expression of *Hsf5* was assessed by reanalyzing scRNA-seq data from adult mouse testes (Hermann et al. 2018) (Fig. S1B). The Uniform Manifold Approximation and Projection (UMAP) of scRNA-seq data suggested that *Hsf5* was coordinately expressed with the landmark genes of meiotic prophase, such as *Spo11*, *Terb1* and *Mybl1* rather than with those of meiotic initiation, such as *Meiosin* and *Stra8,* and those of spermatids during spermiogenesis, such as *Prm1* (Fig. S1B), suggesting that *Hsf5* mRNA was expressed in mouse meiotic prophase. Notably, *Hsf5* showed a different expression pattern from that of other *Hsf* members in mouse testes (Fig. S1B). Although the expression of *Hsf3* and *Hsf4* was barely detected at any spermatogenic stage, *Hsf1, Hsf2,* and *Hsfy2* were expressed in mouse testes. *Hsf1* showed an overall similar expression pattern to *Hsf5* at lower levels. Expression levels of *Hsf2 and Hsfy2* were low at the early meiotic prophase and increased at later spermatogenic stages compared to *Hsf5*. In contrast to spermatogenic expression, *Hsf5* expression was barely detected in E12.5–E18.5 embryonic ovaries by RT-qPCR (Fig.1E), suggesting specific expression of *Hsf5* in male meiotic prophase. These observations suggest that HSF family paralogs may function at different stages in the testis and that HSF5 may play a meiosis-specific role during spermatogenesis.

To determine the stage when HSF5 protein is specifically expressed, seminiferous tubules of the WT mouse testes (8 weeks old) were immunostained with specific antibodies against HSF5 along with SYCP3 (a component of meiotic axial elements) and γH2AX (a marker of DSBs and XY body) or STRA8 (a marker of meiotic initiation) (Fig.1F). The HSF5 signal began to appear in the middle pachytene spermatocyte nuclei of stage VI seminiferous tubules and persisted in the round spermatids of stage VI seminiferous tubules. However, this was not observed in spermatogonia or spermatocytes before mid-pachytene or in elongated spermatids (Fig.1F). Testis-specific histone H1t is a marker of spermatocytes later than mid-pachytene and round spermatids (Drabent et al. 1996) (Cobb et al. 1999). Immunostaining of seminiferous tubules at P16 by H1t along with HSF5 indicated that the HSF5 signal started to appear following the expression of H1t (Fig.1G), confirming the observation above. Although the expression of *Hsf5* mRNA was upregulated upon meiotic entry, immunostaining of the HSF5 protein detected no more than background levels in spermatocytes before the pachytene stage, suggesting that the expression of HSF5 may be post-transcriptionally regulated after entry into the meiotic prophase. These observations suggest that HSF5 is involved in developmental regulation at the mid-to late-pachytene stage of meiotic prophase in males.

### Spermatogenesis was impaired in *Hsf5* knockout males

To address the role of HSF5 in mice, we deleted all coding exons (Exon1-Exon6) of the *Hsf5* loci in C57BL/6 fertilized eggs using the CRISPR/Cas9 system (Fig.2A). Immunoblotting of the extract from *Hsf5* knockout (KO) testes showed that the HSF5 protein was absent (Fig. 2B), which was further confirmed by the diminished immunolocalization of HSF5 in the seminiferous tubules of *Hsf5* KO mice (Fig. 2C), indicating that the targeted *Hsf5* allele was knocked out.

**Figure 2.**
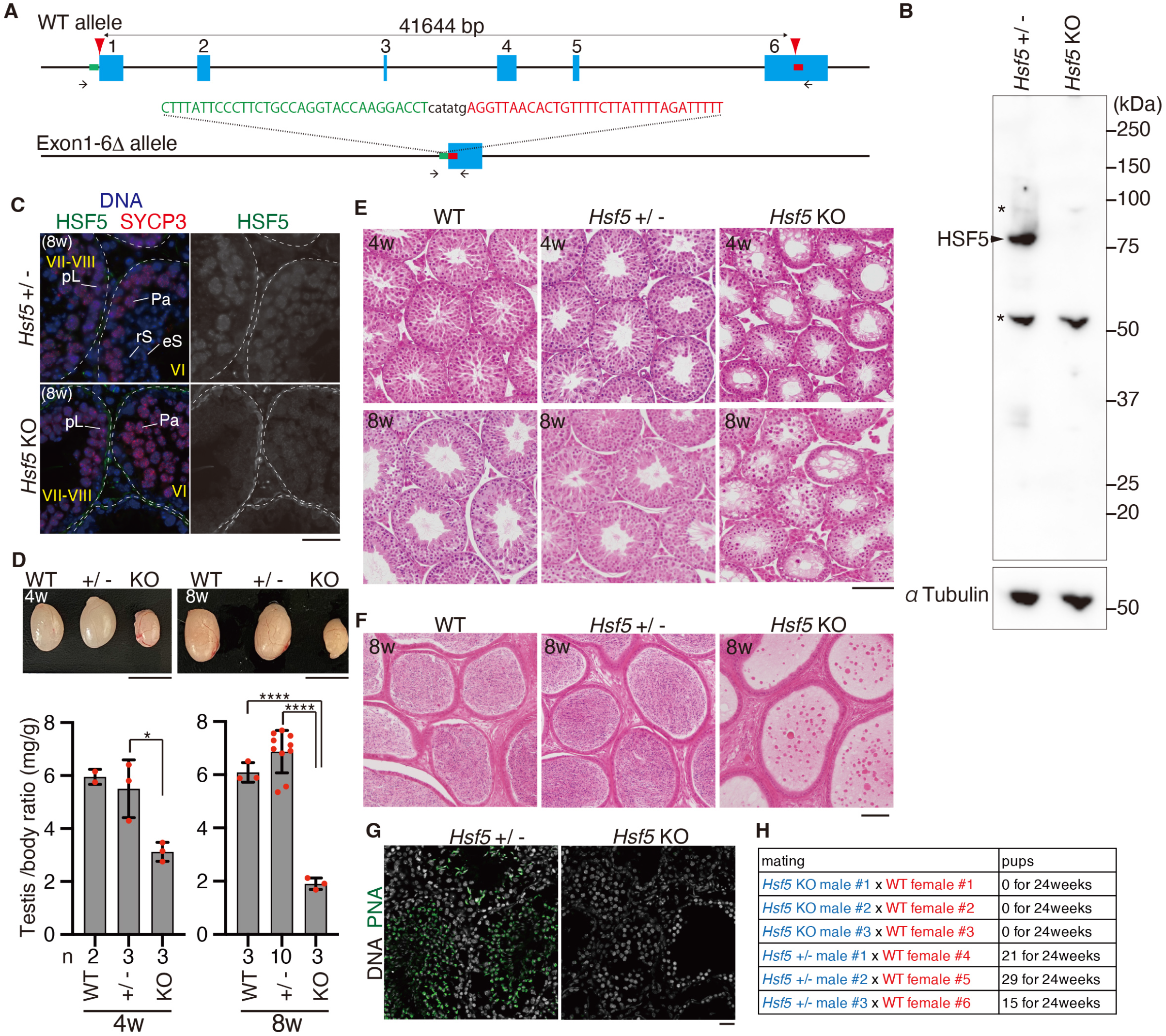
Spermatogenesis was impaired in *Hsf5* knockout male. **(A)** The allele with targeted deletion of Exon1-6 in *Hsf5* gene was generated by the introduction of CAS9, the synthetic gRNAs designed to target upstream of Exon1 and the downstream of Exon6 (arrowheads), and ssODN into C57BL/6 fertilized eggs. 5’- and 3’-homology sequences are shown in green and red, respectively. Arrows: PCR primers for genotyping. Three lines of KO mice (#5, #9, #21) were established. Line #21 of *Hsf5* KO mice was used in most of the experiments, unless otherwise stated. **(B)** Immunoblot analysis of testis extracts prepared from mice with the indicated genotypes (P17). Arrow indicates a band of HSF5. * indicates nonspecific bands. **(C)** Seminiferous tubule sections (8-weeks old) were stained for SYCP3, HSF5 and DAPI. pL: preleptotene, Pa: pachytene spermatocyte, rS: round spermatid, eS: elongated spermatid. Boundaries of the seminiferous tubules are indicated by white dashed lines. Roman numbers indicate the seminiferous tubule stages. Biologically independent mice (N=3) for each genotype were examined. Scale bars: 25 μm. **(D)** Testes from WT, *Hsf5* +/- and *Hsf5* KO (upper left : 4-weeks old, upper right : 8-weeks old left). Testis/body-weight ratio (mg/g) of WT, *Hsf5* +/- and *Hsf5* KO mice (lower left : 4-weeks old, lower right : 8-weeks old) are shown below (Mean with SD). n: the number of animals examined. Statistical significance is shown by *: *p* < 0.01, ****: *p* < 0.0001 (Two-tailed t-test). Scale bar: 5 mm. **(E)** Hematoxylin and eosin staining of the sections from WT, *Hsf5* +/- and *Hsf5* KO testes (upper: 4-weeks old, lower : 8-weeks old). Biologically independent mice (N=3) for each genotype were examined. Scale bar: 100 μm. **(F)** Hematoxylin and eosin staining of the sections from WT, *Hsf5* +/- and *Hsf5* KO epididymis (8-weeks old). Biologically independent mice (N=3) for each genotype were examined. Scale bar: 100 μm. **(G)** Seminiferous tubule sections (8-weeks old) were stained for PNA lectin and DAPI. Scale bar: 25 μm. **(H)** Number of pups born by mating *Hsf5*+/- and *Hsf5* KO males with WT females to examine fertility. *Hsf5* +/- males and *Hsf5* KO males were initially mated with WT females (all 4-weeks old at the start point of mating). This cage was observed for 24weeks from the start of mating.

Although *Hsf5* KO male mice did not show overt phenotypes in somatic tissues, examination of the reproductive organs revealed smaller testes in *Hsf5* KO mice compared to those in WT and *Hsf5* heterozygous mice during the juvenile (4-weeks old) and adult (8-weeks old) periods (Fig. 2D). Histological analysis revealed that post-meiotic spermatids and spermatozoa were absent in 4-weeks and 8-weeks old *Hsf5* KO mice, in contrast to their WT and heterozygous siblings (Fig. 2E). Accordingly, sperm were absent from the adult *Hsf5* KO caudal epididymis at 8 weeks (Fig. 2F). Consistently, seminiferous tubules containing PNA lectin (a marker of spermatids)-positive cells were absent in *Hsf5* KO mice (Fig. 2G). Thus, the later stages of spermatogenesis were severely abolished in *Hsf5* KO seminiferous tubules, resulting in male infertility (Fig. 2H). In contrast to males, *Hsf5* KO females exhibited seemingly normal fertility with no apparent defects in the adult ovaries (Fig. S2A, S2B, S2C). These results suggest that HSF5 is essential for spermatogenesis but not oogenesis.

### HSF5 is required for progression through pachytene in male meiotic prophase

To identify the stage at which the primary defect appeared in the *Hsf5* KO, we analyzed the progression of spermatogenesis by immunostaining seminiferous tubules (4-weeks old) with antibodies against SYCP3 and H1t (a marker of spermatocytes later than mid-pachytene and round spermatids). While all the seminiferous tubules in WT contained H1t-positive spermatocytes and/or round spermatids at 4-weeks old, H1t-positive spermatocytes, but not round spermatids, were observed in around 55% (n = 3) of total seminiferous tubules in the same aged *Hsf5* KO (Fig. 3A). In *Hsf5* KO seminiferous tubules, fewer H1t-positive spermatocytes, if any, were observed and were accompanied by an aberrant staining pattern of SYCP3. These observations suggest that *Hsf5* KO spermatocytes reached at least the mid-pachytene stage, but the progression of meiotic prophase beyond pachytene was seemingly compromised in the absence of HSF5.

**Figure 3.**
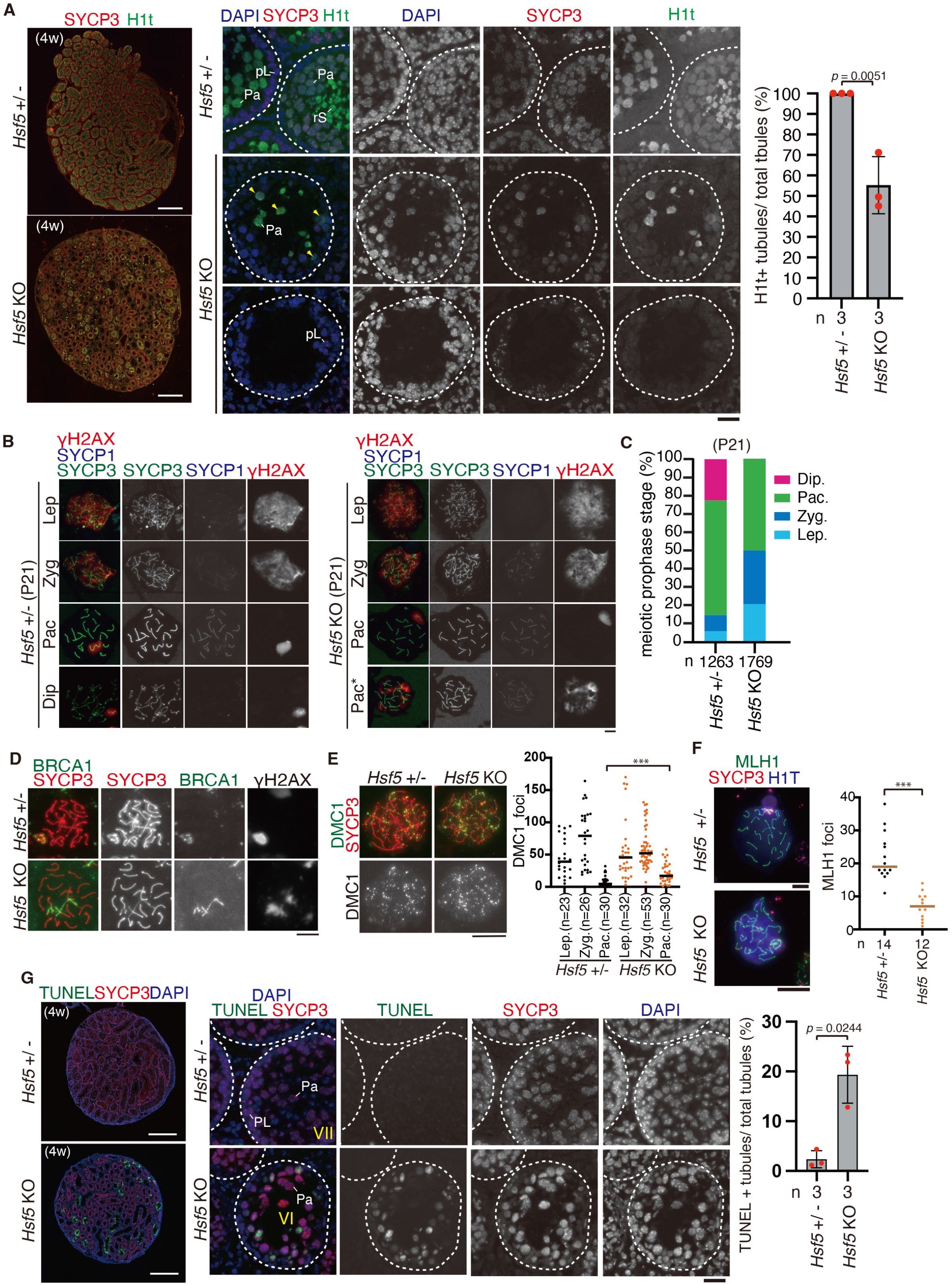
*Hsf5* KO spermatocytes failed to progress through pachytene. **(A)** Seminiferous tubule sections (4-weeks old) were stained for SYCP3, H1t and DAPI. Whole testis sections are shown on the left. Scale bar: 500 μm. Shown on the middle are closeup view of H1t-positive and negative seminiferous tubule sections. Scale bar: 25 μm. pL: preleptotene, Pa: pachytene spermatocyte, rS: round spermatid. Yellow arrowhead indicates H1t-positive spermatocyte with abnormal SYCP3 staining. Shown on the right is the quantification of the seminiferous tubules that have H1t+/SYCP3+ cells per the seminiferous tubules that have SYCP3+ spermatocyte cells in *Hsf5* +/- and *Hsf5* KO testes (Mean with SD). n: the number of animals examined for each genotype. Statistical significance is shown (p = 0.0051, unpaired two-tailed t-test). **(B)** Chromosome spreads of *Hsf5* +/- and *Hsf5* KO spermatocytes at postnatal day 21 (P21) were immunostained as indicated. Lep: leptotene, Zyg: zygotene, Pac: pachytene, Dip: diplotene. Pac* indicates pachytene spermatocyte with high level of γH2AX signals remained on autosomes. Scale bar: 10 μm. **(C)** Quantification of meiotic prophase stage spermatocytes per total SYCP3+ spermatocytes in *Hsf5* +/- and *Hsf5* KO mice at P21. n: the number of cells examined. **(D)** Chromosome spreads of *Hsf5* +/- and *Hsf5* KO pachytene spermatocytes were stained for BRCA1, SYCP3, and γH2AX. ∼17.8 % of *Hsf5* KO pachytene spermatocytes (N=62) exhibited BRCA1 along autosomes with γH2AX signals, whereas none of *Hsf5* +/- pachytene spermatocytes (N=51) did except for XY chromosome. Scale bar: 10 μm. **(E)** Chromosome spreads of *Hsf5* +/- and *Hsf5* KO spermatocytes were stained for DMC1, SYCP3, and SYCP1. Immunostained chromosome spread of zygotene spermatocytes are shown. The number of DMC1 foci is shown in the scatter plot with median (right). Statistical significance is shown by *p*-value (Mann-Whitney U-test). ***: p < 0.0001. Lep.: leptotene, Zyg.: Zygotene, Pac.: Pachytene. Scale bar: 10 μm. **(F)** Chromosome spreads of *Hsf5* +/- and *Hsf5* KO spermatocytes were stained for MLH1, SYCP3 and SYCP1. The number of MLH1 foci is shown in the scatter plot with median (right). Statistical significance is shown (Mann-Whitney U-test). n: number of spermatocytes examined. Statistical significance is shown by *p*-value (Mann-Whitney U-test). ***: p < 0.0001. Scale bar: 5 μm. **(G)** Seminiferous tubule sections from 4-weeks old mice were subjected to TUNEL assay with immunostaining for SYCP3. Whole testis sections are shown on the left. Scale bar: 500 μm. Shown on the middle are closeup view of TUNEL-positive and negative seminiferous tubule sections. Scale bar: 25 μm. pL: preleptotene, Pa: pachytene. Shown on the right is the quantification of the seminiferous tubules that have TUNEL+ cells per total tubules in *Hsf5* +/- (n=3) and *Hsf5* KO (n=3) testes (bar graph with SD). Statistical significance is shown by *p*-value (*p* = 0.0244, unpaired two-tailed t-test).

Immunostaining analysis with antibodies against SYCP3 and SYCP1 (markers of homolog synapsis) and γH2AX demonstrated that spermatocytes underwent homologous chromosome synapsis in juvenile *Hsf5* KO males (P21), as in age-matched controls (Fig 3B). However, spermatocytes after pachytene and post-meiotic spermatids were not observed in *Hsf5* KO mice at P21 (Fig. 3C). Accordingly, more leptotene/zygotene and reciprocally fewer pachytene populations were observed in *Hsf5* KO spermatocytes than in age-matched controls (Fig. 3C), suggesting that *Hsf5* KO spermatocytes were arrested at pachytene.

Normally, the first wave of γH2AX is mediated by ATM after double-strand break (DSB) formation in leptotene (Mahadevaiah et al. 2001) and disappears during DSB repair. The second wave of γH2A in the zygotene stage is mediated by ATR, which targets unsynapsed or unrepaired chromosomes (Royo et al. 2013). At the leptotene and zygotene stages, γH2AX signals appeared in *Hsf5* KO spermatocytes in the same manner as in WT (Fig. 3B), indicating that DSB formation occurred normally in *Hsf5* KO spermatocytes. However, atypical γH2AX staining patterns in *Hsf5* KO pachytene spermatocytes were observed. Specifically, γH2AX signals largely persisted throughout the nuclei, including on fully synapsed autosomes, in ∼24.2% of *Hsf5* KO pachytene spermatocytes (N=62), whereas they disappeared in the control pachytene spermatocytes for retaining on the XY body. Furthermore, BRCA1, a marker of DNA damage response (Turner et al. 2004) (Broering et al. 2014), appeared along autosomes in ∼17.8% of *Hsf5* KO pachytene spermatocytes (N=62) (Fig 3D). These observations suggested that DSBs were not repaired and/or were newly generated in *Hsf5* KO spermatocytes. Consistently, the number of DMC1 foci (a marker of ssDNA at the DBS site) was significantly increased in *Hsf5* KO pachytene spermatocytes, suggesting that DSBs were yet to be fully repaired in some, if not all, *Hsf5* KO pachytene spermatocytes (Fig 3E). Accordingly, the number of MLH1 foci (a marker of crossover recombination) was reduced in *Hsf5* KO pachytene spermatocytes compared to that in the control (Fig. 3F), suggesting that crossover recombination was incomplete in *Hsf5* KO pachytene spermatocytes. These results suggest that DBS repair and crossover formation are defective in *Hsf5* KO pachytene spermatocytes despite fully synapsed homologs.

Notably, a higher number of TUNEL-positive seminiferous tubules (∼19.3% of total tubules) was observed in *Hsf5* KO testes at 4 weeks (Fig. 3G). Since TUNEL-positive cells were observed in stage VI seminiferous tubules that contained mid-pachytene spermatocytes in *Hsf5* KO mice (Fig. 3G), ongoing germ cell degeneration presumably occurred at mid-pachytene in *Hsf5* KO spermatocytes, at the time when HSF5 first appeared during spermatogenesis (Fig.1F). These observations suggest that *Hsf5* KO spermatocytes failed to progress through the pachytene stage and were consequently eliminated by apoptosis. Therefore, HSF5 is required for progression through the pachytene stage of the meiotic prophase.

### *Hsf5* KO spermatocytes showed alteration of gene expression at meiotic prophase

HSF1 and HSF2 are implicated in spermatogenesis under non-stress conditions (Salmand et al. 2008) (Kallio et al. 2002) (Wang et al. 2003) (Wang et al. 2004). Given that *Hsf5* KO spermatocytes failed to progress beyond the pachytene stage under non-stress conditions in our cytological analyses (Fig.3), we conducted transcriptome analysis to determine whether *Hsf5* KO spermatocytes had altered gene expression profiles under non-stress conditions. For this purpose, we isolated spermatocytes that were in the progression of meiotic prophase by fluorescent sorting with DyeCycle Violet (DCV) staining from WT and *Hsf5* KO testes (Fig. S3)(Yeh et al. 2021). Since H1t-negative early pachytene is the stage before defects appeared in the mutants (Fig.3A), we assumed that the cellular composition should be similar in the control WT and *Hsf5* KO until the first wave of meiotic prophase reaches the mid to late pachytene stage. This allowed for the comparison of the transcriptomes of the sorted cells in WT and *Hsf5* KO mice with minimized batch effects that could potentially be caused by a bias in the cellular population. Since the number of sorted populations was limited, we conducted SMART RNA-seq on the sorted spermatocytes, which allowed for RNA-seq analysis with small cell numbers (Fig.4).

**Figure 4.**
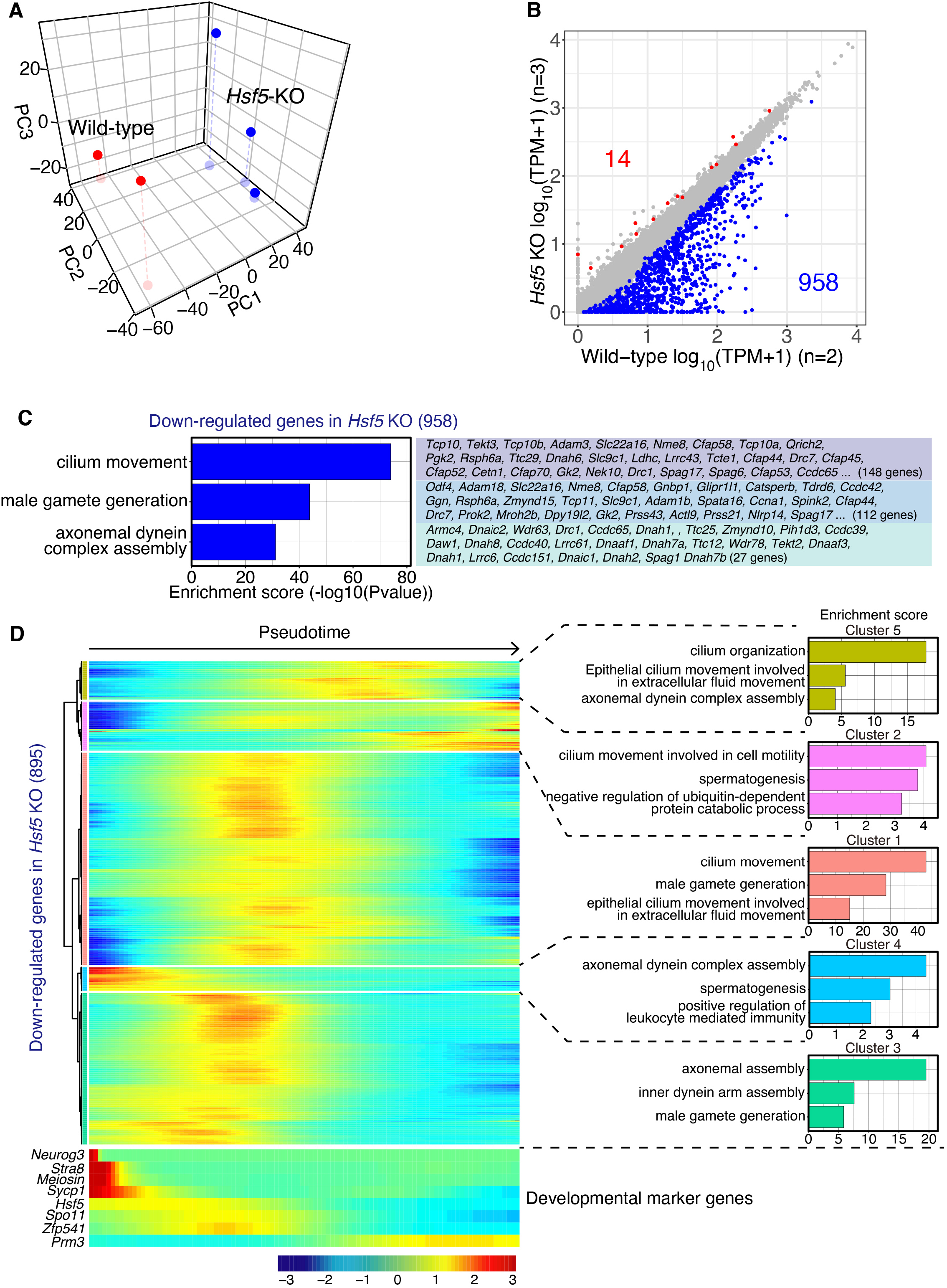
SMART-seq analysis of meiotic prophase spermatocytes in *Hsf5* KO. **(A)** The early meiotic prophase spermatocytes (zygotene/pachytene) were isolated from control WT (n=3) and *Hsf5* KO (n=2) testes at P17 by fluorescent sorting, and subjected to SMART RNA-seq. Principal component analysis of the transcriptomes of meiotic prophase spermatocytes in WT and *Hsf5* KO is shown. **(B)** Scatter plot of the transcriptome of meiotic prophase spermatocytes in WT versus *Hsf5* KO is shown. The numbers of differentially expressed genes are shown. Significance criteria: false discovery rate ≤ 0.05. **(C)** Gene enrichment analysis of the 958 downregulated genes in meiotic prophase spermatocytes of *Hsf5* KO testes. The enriched term for 14 upregulated genes is not shown due to low statistical significance. See Supplementary Data 2 for complete gene list of the Gene enrichment analyses. **(D)** Heatmap showing the hierarchical relationship among the clusters of the downregulated genes in *Hsf5* KO across pseudotime of spermatogenesis. Expressions of the downregulated genes in meiotic prophase spermatocytes of *Hsf5* KO was assessed by reanalyzing scRNA-seq data of spermatogenic cells (GEO : GSE109033) (Hermann et al. 2018). Pseudotime (left to right) corresponds to developmental trajectory of spermatogenesis (undifferentiated spermatogonia to round spermatids). For a reference, Expression profiles of key developmental marker genes are shown along pseudotime. The enriched terms (top 3 by enriched score) are shown on the right. See Supplementary Data 2 for the complete gene list of the Gene enrichment analyses.

Principal component analysis (PCA) revealed that the overall transcriptomes of enriched spermatocytes in *Hsf5* KO testes differed from those in WT testes (Fig.4A). Among the differentially expressed genes (DEG) between WT and *Hsf5* KO pachytene spermatocytes, 958 genes were downregulated in *Hsf5* KO mice, whereas 14 genes were upregulated in *Hsf5* KO mice (Fig.4B). Notably, gene enrichment analysis indicated that the genes involved in cilium movement, male gamete generation, and sperm axonemal dynein complex assembly were downregulated in *Hsf5* KO mice (Fig. 4C, Supplementary Data 2). In contrast, the genes upregulated in *Hsf5* KO mice (14 genes) were not significantly associated with any Gene enriched terms. Reanalysis of the downregulated genes in *Hsf5* KO (958 genes) using previously published scRNA-seq data from spermatogenic cells (Hermann et al. 2018) suggested that these genes were expressed around the mid-stage of pseudotime and declined after that (Fig. 4D). Thus, these results suggest that the gene expression pattern was different between WT and *Hsf5* KO spermatocytes during meiotic prophase, although it is still possible that subtle differences in developmental cellular populations between control WT and *Hsf5* KO spermatocytes may have potentially contributed to bulk transcriptomic differences.

### Failure of developmental progression beyond a substage of mid-pachytene in *Hsf5* KO spermatocytes

Since the pachytene stage is solely defined by the cytologically defined appearance of chromosomal morphology (spermatocytes with fully synapsed homologs) and covers a broad developmental period, it remains unclear whether a specific subtype of pachytene spermatocytes was eliminated during spermatogenesis, or whether a newly emerged subtype caused the primary defect in *Hsf5* KO mice. To identify which subtypes of *Hsf5* KO spermatocytes accompanied the alteration of gene expression profiles, we further conducted scRNA-seq analyses of whole testicular cells from WT and *Hsf5* KO mice at postnatal day 16 (Fig. S4). Although the cellular composition in the testes changes with developmental progression, we assumed that the first wave of spermatocytes would progress with the same cellular composition in WT and *Hsf5* KO mice until mid-pachytene. We sought a subtype of spermatocytes that would be present in WT but not in *Hsf5* KO testes, or vice versa, predicted from gene expression patterns at this time point, because it would approximately cover the transition when HSF5 started to exhibit its mandatory function in WT spermatocytes and when the primary defect appeared in *Hsf5* KO spermatocytes. Since HSF5 expression was restricted to germ cells, scRNA-seq datasets derived from germ cell populations (spermatogonia and spermatocytes) were analyzed separately from those of testicular somatic cells (Sertoli cells, Leydig cells, peritubular myoids, endothelial cells, and hemocytes) (Fig. 5A, Fig. S4A, S4B). The UMAP of the scRNA-seq dataset indicated that the gene expression patterns of single cells isolated from WT and *Hsf5* KO spermatogenic germ cell populations were separated into 12 clusters (Fig. 5B, C, Fig. S4C, S4D, and Supplementary Data 3). The expression patterns of key marker genes were used to estimate the developmental direction of spermatogenesis at the P16 on the UMAP (Fig. 5D). Cluster 11 represented the SSC population, as suggested by the high expression *Gfra1* and *Zbtb16*. Cluster 8 represented an undifferentiated spermatogonial population, as suggested by the high expression *Zbtb16*. Cluster 7 represented the population of differentiating spermatogonia, as suggested by the upregulation of *Kit.* Cluster 1 represents the population at meiotic initiation, as suggested by the upregulation of *Meiosin.* Clusters 0 and 9 represented the early meiotic prophase population, as suggested by the upregulation of *Spo11.* Clusters 3, 2, and 10 represented the mid-meiotic prophase populations, as suggested by the upregulation of *Tesmin.* Based on these, we assumed that spermatogenesis progressed along the trajectory from Cluster 11 to Cluster 10 in the UMAP.

**Figure 5.**
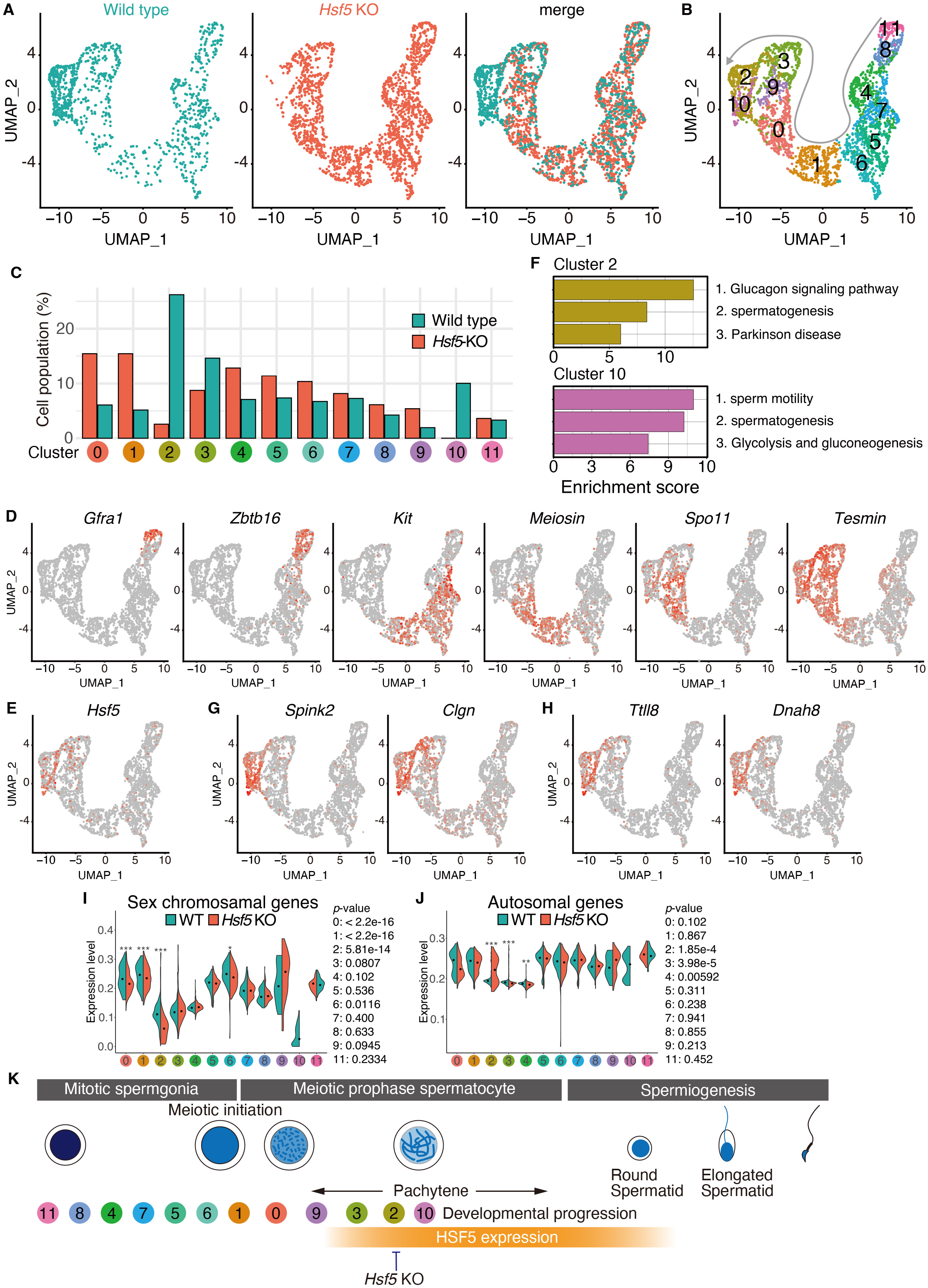
scRNA-seq analysis of WT and *Hsf5* KO spermatogenic germ cells. **(A)** UMAP representation of scRNA-seq transcriptome profiles for germ cells from P16 WT and *Hsf5* KO testes. **(B)** Clustering analysis of different gene expression patterns on UMAP-defined scRNA-seq transcriptomes of P16 WT and *Hsf5* KO cells. Gray arrow indicates developmental direction. **(C)** Bar graph showing the proportion of WT and *Hsf5* KO germ cells among the clusters. **(D)** UMAP plots show expression patterns of key developmental marker genes of spermatogenic cells. Key developmental marker genes include *Gfra1* : spermatogonial stem cell, *Zbtb16* : undifferentiated spermatogonia, *Kit* : differentiating spermatogonia, *Meiosin*: pleleptotene spermatocyte, *Spo11*: early meiotic prophase spermatocyte, *Tesmin*: mid-pachytene spermatocyte. **(E)** Expression pattern of *Hsf5* on the UMAP plot. **(F)** Gene enrichment analysis of highly expressed genes in Cluster 2 or Cluster 10. **(G)** Expression patterns of the representative genes in Cluster 2 (*Spink2, Clgn*) on the UMAP plot. **(H)** Expression patterns of the representative genes in Cluster 10 (*Ttll8, Dnah8*) on the UMAP plot. **(I)** Expression levels of the sex chromosomal genes among the clusters are shown in violin plots with a median. *p*-values by Wilcoxon rank sum test are shown on the right. **(J)** Expression levels of the autosomal genes among the clusters are shown in violin plots with a median. *p*-values by Wilcoxon rank sum test are shown on the right. **(K)** The subtype clusters delineated by scRNA-seq and the timing of HSF5 protein expressions are shown along the developmental stages. HSF5 started to appear in the spermatocyte nuclei from mid-pachytene onward, and were expressed in round spermatids. Vertical bars indicate the stages when the developmental progression is blocked in *Hsf5*KO spermatocytes. See also Figure S4, Supplementary Data 3.

The gene expression profiles of WT and *Hsf5* KO germ cell populations were well overlapped along the trajectory from Clusters 11 to 3. However, we noticed subpopulations (Clusters 2 and 10) that were present in WT but largely missing in *Hsf5* KO testes (Fig. 5B, C), presumably corresponding to the most advanced stages of meiotic prophase in WT spermatocytes at P16. Intriguingly, Clusters 2 and 10 coincided with the abrupt upregulation of *Hsf5* expression in the WT cells (Fig. 5E). Thus, the subtypes of spermatocytes that represented Clusters 2 and 10 were absent in *Hsf5* KO spermatocytes, which were reciprocal to the upregulation of *Hsf5* in WT testes in Clusters 2 and 10. Consistent with the cytological observation that *Hsf5* KO spermatocytes progressed through the early meiotic prophase but were eliminated by apoptosis at mid-pachytene (Fig. 3), HSF5 was required for spermatocytes to progress beyond the substages of meiotic prophase that corresponded to Clusters 2 and 10.

Gene enrichment analysis revealed that genes related to spermatogenesis (*Spink2, Clgn, Hspa2, Ybx3, Tcp11, Rsph1, Pebp1, Ybx2, Nphp1, Catsperz, Ggnbp1, Psma8, Meig1, Ropn1l, Morn2*) were highly expressed in Clusters 2 and 10 (Fig. 5F, G, Supplementary Data 3). Furthermore, genes related to sperm motility, such as (*Ttll8, Dnah8, Ldhc, Sord, Ccdc39, Gk2, Cfap206, Zmynd10, Ropn1l, Spata33, Dnaaf1*) were highly expressed in Cluster 10 (Fig. 5F, H, Supplementary Data 3). Since *Spink2* and *Clgn* are known to be expressed in pachytene spermatocytes, Clusters 2 and 10 represent the subtypes of pachytene spermatocytes. In contrast, genes associated with Clusters 2 and 10 were underrepresented in *Hsf5* KO spermatocytes. Crucially, Cluster 10 coincided with the abrupt downregulation of the sex chromosome genes (Fig. 5I), suggesting that those Cluster 2 represents the subpopulation of mid-pachytene when meiotic sex chromosome inactivation (MSCI) established. Note that *Hsf5* KO spermatocytes exhibited downregulation of sex chromosome genes in Cluster 2, suggesting that *Hsf5* KO spermatocytes at least exhibited a sign of undergoing MSCI at this time point but were yet to establish MSCI. Notably, overall expression level of autosomal genes was abruptly upregulated in Cluster 2 subpopulation of *Hsf5* KO (Fig. 5J), suggesting that some of, if not all, autosomal genes were derepressed in the absence of HSF5 in the subpopulation of Clusters 2. Thus, although *Hsf5* KO spermatocytes exhibited gene expression patterns that were comparable to those in WT spermatocytes before the mid-pachytene, specific subpopulations of mid-pachytene spermatocytes represented by Clusters 2 and 10 were lost in *Hsf5* KO testes (Fig. 5K). Therefore, HSF5 is essential for the developmental progression beyond the mid-pachytene substage during spermatogenesis.

### HSF5 binds to the promoter regions of the genes with a unique target specificity

HSF5 was predicted to possess a putative DNA-binding domain (Fig. 1B). We further investigated HSF5-target sites in the genome using ChIP-seq analysis. HSF5 bound to 165 sites (161 nearest genes, which were assigned regardless of the distance from the HSF5-binding sites), of which 93.9% resided within 1kb of the transcription start sites (TSS) in the promoter regions of the mouse genome (Fig. 6A-C, Supplementary Data 4). Notably, DNA-binding motif analysis identified *de novo* HSF5-ChIP enriched DNA sequences that were different from the previously known binding motifs of other HSF family transcription factors (Fig. 6D). While the typical HSE that resides in the promoter regions of heat shock-responsive genes is composed of three contiguous inverted repeats, nTTCnnGAAnnTTCn (Trinklein et al. 2004), the most probable HSF5-binding motif is composed of the octamer sequence (T/C/G)(G/A)GAA(C/T)(G/T/C)(C/T) with a strong preference for a single triplet GAA (Fig. 6D). This suggests that HSF5 directly binds to promoters through this motif, which is distinct from HSE bound by other canonical HSF family transcription factors. It should be noted that HSF5-target genes were little overlapped with either of HSF1-target genes (Akerfelt et al. 2010b) or HSF2-target genes in the testis (Akerfelt et al. 2008) (Fig. 6E, Supplementary Data 4). Thus, HSF5 binds to different target genes to those bound by HSF1 or HSF2 during spermatogenesis.

**Figure 6.**
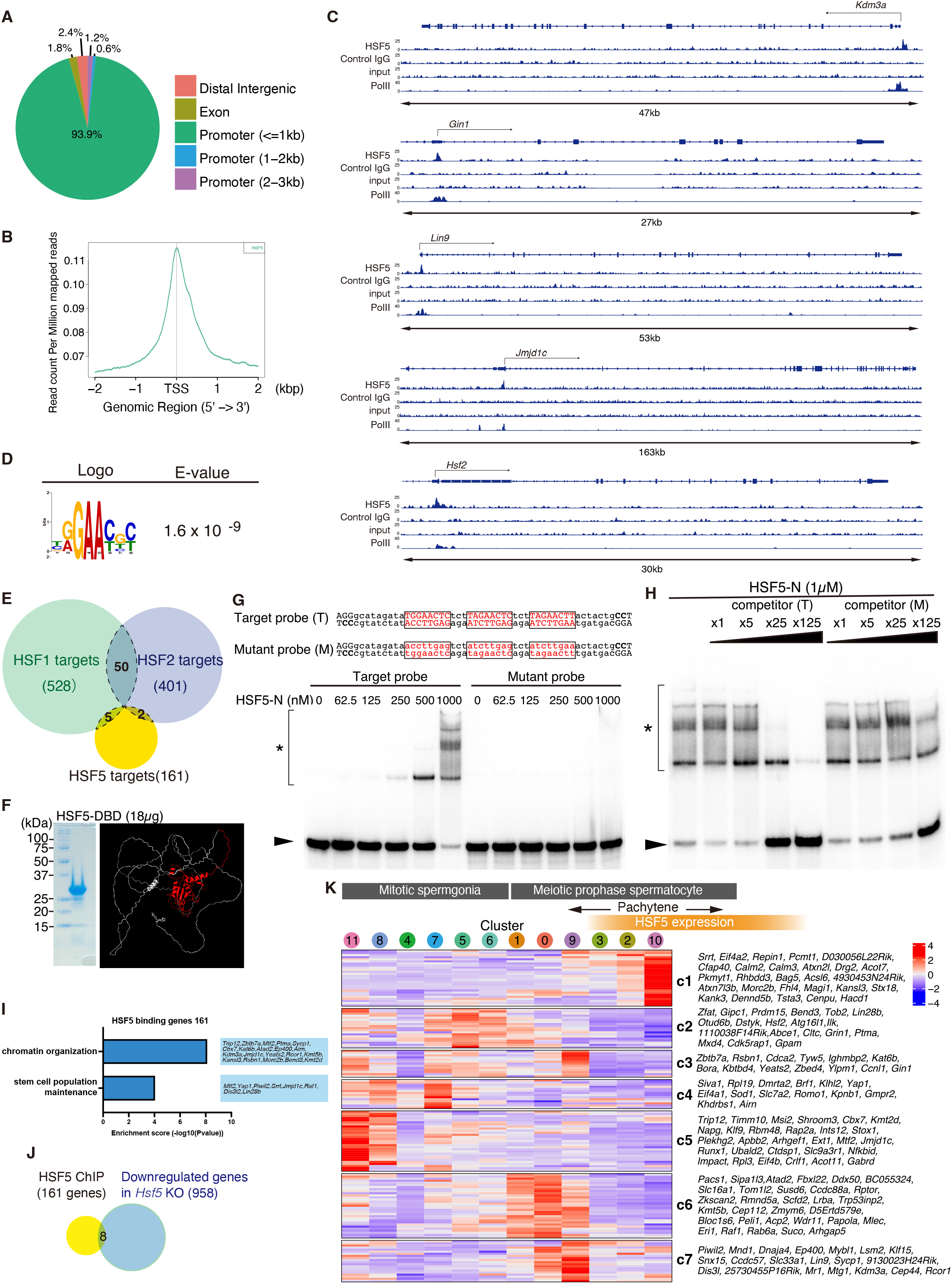
HSF5 binds to the gene promoter regions. **(A)** HSF5 binding sites were classified by the genomic locations as indicated. **(B)** Heat map of the common HSF5 binding sites of HSF5 ChIP-seq at the positions - 2.0 kb upstream to +2.0 kb downstream relative to the TSS. Average distributions of HSF5-ChIP-seq binding peak are shown on the bottom. **(C)** Genomic view of HSF5 ChIP-seq control IgG ChIP-seq input DNA data over representative gene loci. Genomic coordinates were obtained from RefSeq. RefSeq IDs for mRNA isoforms are indicated. To specify testis specific transcription, RNA polymerase II ChIP-seq in the testis are shown (Li et al. 2013). **(D)** The top sequence motif enriched in HSF5 ChIP-seq with E-values. MEME-ChIP E-value estimates the expected number of motifs with similar features that one would find in a similarly sized set of random sequences. See also Supplementary Data 4. **(E)** Venn diagram representing the overlap among the targets of HSF1 (Akerfelt et al. 2010b), HSF2 (Akerfelt et al. 2008), and HSF5 in the testes. **(F)** The purified HSF5 N-terminal (aa1-209) protein was used for EMSA assay (left). Shown on the right is the corresponding HSF5 N-terminal part is shown (red) on the ribbon model that was predicted from AlphaFold2. **(G)** DNA binding ability of HSF5 DNA binding domain was examined by EMSA assay. Shown on the top are the target (T) and mutant (M) sequences of the DNA probes. Target (T) sequence was designed according to the enriched motif that was predicted by Chip-seq as shown in (D). Increasing amount of the purified protein was mixed with 0.04 pmol of ^32^P-labeled DNA probes (T or M) at the protein/DNA molar ratio of 15.6 - 250. Arrowhead: unbound DNA. The protein-DNA complexes are shown by * with a bracket. **(H)** HSF5 N-terminal protein (1μM) was mixed with 0.04 pmol of the ^32^P-labeled target (T) DNA probe. DNA binding specificity of HSF5-DNA complex was assessed by adding the unlabeled target or mutant competitor DNA (1 – 125 fold excess to the ^32^P-labeled DNA probes). **(I)** Gene enrichment analysis of the HSF5-bound genes (161genes). Top 2 biological processes ranked by log (*p*-value) are listed. See Supplementary Data 4 for complete list of the HSF5-bound genes and the Gene enrichment analyses. We used the standard parameters to detect over-represented enriched terms for biological function, giving Fisher’s exact *p*-values. **(J)** Venn diagram representing the overlap of HSF5-bound genes (161 nearest genes) and downregulated (958 genes) in *Hsf5* KO mice. None of HSF5-bound genes overlapped with the upregulated genes (14 genes) in *Hsf5* KO mice. **(K)** Heatmaps showing the hierarchical relationship of the expression patterns of the HSF5-bound genes across the developmental direction. Expressions of the HSF5-bound genes were assessed by scRNA-seq data of spermatogenic cells as described in Fig.5I. The cluster number is indicated on the bottom. The order of clusters from left to right corresponds to developmental direction of spermatogenesis (undifferentiated spermatogonia to pachytene spermatocyte). The expression patterns of the HSF5-bound genes were classified (c1-c7). See also Supplementary Data 4.

To validate the DNA-binding specificity of HSF5 to the motif predicted from the HSF5 ChIP-seq data, we examined the DNA-binding ability of the HSF5 DNA-binding domain (HSF5-DBD) using an *in vitro* EMSA (Fig. 6F-H). Indeed, HSF5-DBD bound to DNA with the target motif but not DNA with mutant sequences (Fig. 6G). As the probe contained three octamer motif units, three shifted bands appeared when increasing amounts of HSF5-DBD were added. In contrast, all mobility shifts of the bands were completely abolished when substitutions were introduced into the octamer motif. We confirmed that the HSF5 protein-DNA complex was titrated away in the presence of an excess amount of unlabeled DNA with the target motif but not the mutant sequence, showing the DNA-binding specificity of HSF5 to the predicted motif (Fig. 6H). Thus, the predicted motif with a single triplet GAA was sufficient for the DNA binding of HSF5-DBD.

Remarkably, Gene enrichment analysis revealed that the putative HSF5-bound genes were associated with the biological processes of chromatin organization and stem cell population maintenance (Fig. 6I, Supplementary Data 4). It should be noted that most of these genes are generally expressed in broad cell types rather than being germ cell-specific. Among the putative HSF5-bound genes, only eight and none were identified among the down- and up-regulated genes in *Hsf5* KO meiotic prophase spermatocytes, respectively (Fig. 6J). This is presumably because *Hsf5* KO spermatocytes were eliminated soon after they reached a specific point in the pachytene stage when the HSF5 protein was upregulated (Fig. 5). This hampered *bona fide* comparison of the transcriptomes of WT and *Hsf5* KO spermatocytes at a specific time point when HSF5 acts for transcription.

Additionally, we analyzed the expression patterns of HSF5-bound genes during spermatogenesis using the scRNA-seq data from spermatogenic cells (Fig. 5). Hierarchical clustering revealed the expression patterns of HSF5-bound genes in single cells across the order of the scRNA-seq Clusters that corresponded to the developmental direction (Fig. 6K, Supplementary Data 4). The expression levels of HSF5-bound genes were higher in spermatogonia (Clusters 11, 8, 4, 7, 5, and 6 populations in scRNA-seq) and spermatocyte populations at earlier time points of meiotic prophase (Clusters 1, 0, and 9) and then declined before Cluster 2 (Fig. 6K). This observation was consistent with the fact that overall expression level of autosomal genes was abruptly derepressed in Cluster 2 subpopulation of *Hsf5* KO (Fig. 5J). The other class of HSF5-bound genes (class 1) showed higher expression levels in Clusters 2 and 10 (Fig. 6K). These observations suggest that HSF5 positively or negatively regulates the expression of its HSF5 target genes.

To elucidate how HSF5 is involved in transcriptional regulation, factors interacting with HSF5 were screened by immunoprecipitation (IP) using different HSF5 antibodies followed by mass spectrometry (MS). Our HSF5 IP-MS analysis demonstrated that HSF5 immunoprecipitated with the subunits of the SWI/SNF chromatin-remodeling complex, SMARCA5, SMARCC2, SMARCE1 and SMARCC1 (Fig. S5A, S5B, Supplementary Data 5), HSF5 may collaborate with the SWI/SNF complex to positively or negatively regulate HSF5 target gene expression. Notably, HSF5 was repeatedly immunoprecipitated with KCTD19, and its interacting proteins ZFP541, and HDAC1 (Fig. S5A, S5B, Supplementary Data 5). Our previous study demonstrated that KCTD19 reciprocally co-immunoprecipitates with HSF5 in chromatin extracts (Horisawa-Takada et al. 2021). KCTD19 forms a subcomplex that consists of KCTD19, HDAC1/2, TRERF1, and TDIF1, and interacts with DNA-binding proteins such as ZFP541 and MIDEAS, thus acting as a corepressor of transcriptional repression during spermatogenesis (Horisawa-Takada et al. 2021) (Oura et al. 2021). Therefore, HSF5 may repress the target genes collaborating with the KCTD19-HDAC1/2 containing complex.

## DISCUSSION

### HSF5 plays a specific role in the developmental progression of spermatocytes under non-stress conditions

In mouse testes, HSF5 and other HSF family members, HSF1, HSF2, and HSFY2, are expressed at different or overlapping stages of spermatogenesis (Fig. S1B) (Sarge et al. 1994). The testis is a heat-sensitive organ where spermatogenesis occurs under low-temperature conditions (Mieusset and Bujan 1995) (Sarge 1995). In mice, when the testes experience temperature elevation, spermatogenesis is compromised during meiotic recombination, leading to the elimination of spermatocytes (Hirano et al. 2022). A previous study showed that transgenic mice constitutively overexpressing an active form of HSF1 in the testes are infertile due to a block in spermatogenesis and apoptosis, whereas female fertility is unaffected (Nakai et al. 2000) (Izu et al. 2004) (Widlak et al. 2003). Thus, HSF1 acts, at least in part, as a stress response factor in the cell-death decision at the pachytene stage under the stress condition in the testes. In contrast, HSF5 plays a specific role in developmental progression under non-stress conditions in mice (Fig 2, 3).

It has been shown that HSF1 and HSF2 play a role in spermatogenesis in non-stress conditions. *Hsf1* KO (Salmand et al. 2008) and *Hsf2* KO (Wang et al. 2003) (Kallio et al. 2002) males exhibit reduced spermatogenesis but are fertile. In *Hsf1* and *Hsf2* double KO males, spermatocytes fail to progress to the pachytene stage, leading to a complete lack of mature sperm, resulting in male infertility (Wang et al. 2004). HSF1 and HSF2 regulate the expression of the common target genes on the sex chromosomes in spermatogenic cells (Akerfelt et al. 2008) (Akerfelt et al. 2010b). Thus, HSF1 and HSF2 have complementary or overlapping roles in meiotic prophase progression, as suggested by their ability to heterodimerize (Sandqvist et al. 2009). Similarly, we showed that *Hsf5* KO spermatocytes failed to progress beyond the pachytene stage and were consequently eliminated by apoptosis (Fig 3G). Whereas HSF1 faintly appears in the nuclei of early pachytene spermatocytes of the stage I, localizing to the sex body in pachytene spermatocytes and the chromocenter in round spermatids (Akerfelt et al. 2010b), HSF5 appears in the nuclei of mid pachytene spermatocyte, and in the round spermatids with rather excluded from the chromocenter (Fig.1F), indicating the spatially and temporally different behaviors of HSF 1 and HSF5. Crucially, these evidences suggest that HSF5 has a specific function in meiotic prophase progression that cannot be compensated for by HSF1 or HSF2 and vice versa. Therefore, HSF5 plays a distinct role in spermatogenesis, different from HSF1 and HSF2 in mice. Consistently, in zebrafish, *Hsf5* mutant males are infertile with arrest at the zygotene-pachytene transition but still show a normal heat stress response (Saju et al. 2018). Therefore, the role of HSF5 in the progression of spermatogenesis under non-stress conditions is evolutionarily conserved in mammals and fish.

HSFY has been implicated in male infertility in humans (Tessari et al. 2004). Although *Hsfy2* is expressed in spermatogenic cells in the mouse testes (Fig. S1B), it is unknown whether mouse HSFY2 is involved in spermatogenesis. Nevertheless, it is unlikely that HSFY2 has an overlapping function that genetically complements the role of HSF5 in *Hsf5* KO pachytene spermatocytes.

### HSF5 acts as an atypical HSF transcription factor

HSF5 differs from other canonical HSFs (HSF1, HSF2, and HSF4) in terms of protein structure (Fig. 1B), expression pattern (Fig. 1D), DNA-binding specificity (Fig. 6), and physiological function (Fig. 2-5). HSF1, HSF2, and HSF4 were ubiquitously expressed in a wide range of tissues, while HSF5 is specifically expressed in spermatogenic cells in the testis (Fig. 1D). HSF family members are classified as transcription factors that possess an evolutionarily conserved winged-helix-turn-helix (wHTH) DNA-binding domain in fungi, invertebrates, and vertebrates, and were originally described to recognize a consensus HSE (Gomez-Pastor et al. 2018).

We showed that HSF5 recognizes a specific DNA motif different from the typical HSE bound by other canonical HSF family transcription factors (Fig 6). While the typical HSE is composed of three contiguous inverted repeats of nGAAn; (nGAAnnTTCnnGAAn) or its complementary sequence (nTTCnnGAAnnTTCn) (Trinklein et al. 2004) (Gomez-Pastor et al. 2018), our ChIP-seq data predicted that HSF5-binding motif is composed of the octamer (T/C/G)(G/A)GAA(C/T)(G/T/C)(C/T) with a strong preference of a single triplet GAA (Fig. 6D). The DBDs of HSF1, HSF2, and HSF4 possess a recognition helix that contains a conserved Ser-Phe-Val-Arg-Gln amino acid sequence inserted into the major groove of the HSE, in which a conserved Arg residue forms hydrogen bonds with the guanine of GAA and is essential for DNA binding of the HSF (Littlefield and Nelson 1999). In contrast, HSF5 possesses a Ser-Phe-Ile-Arg-Gln amino acid sequence at the same position (Fig. S6), in which Ile, with its bulky side chain, was placed instead of Val at the neighboring position of the essential Arg. Strikingly, HSF5 possesses an insertion of an amino acid sequence between helix 2 and the DNA recognition helix, which corresponds to the predicted IDR (Fig. 1B). The wing domain is poorly conserved in HSF5; however, it appears to have been substituted with a predicted IDR (Fig. 1B). Furthermore, whereas HSF1, HSF2, and HSF4 possess two heptad repeats, HR-A and HR-B, that are predicted to form inter-molecular leucine zippers for homotrimer oligomerization (Rabindran et al. 1993), HSF5 lacks these heptad repeats (Fig. 1B). It is possible that because of these structural differences, HSF5 acquires target specificity that is distinct from that of HSF1, HSF2, and HSF4. Indeed, HSF2 binds to and regulates genes on the Y chromosome long arm (MSYq), such as *Sly and Ssty2* in spermatogenic cells (Akerfelt et al. 2008). HSF1 also binds to the multi-copy genes on the sex chromosomes that are shared by HSF2, and other genes (Akerfelt et al. 2010b). Only 2 genes were commonly bound by HSF1 and HSF5, and a gene was commonly bound by HSF2 and HSF5 (Fig. 6E, Supplementary Data 4). These evidences highlight the different target specificities of HSF1, HSF2 and HSF5 during spermatogenesis. Since HSF5 is theoretically assumed to find the octamer motif at the probability of one binding site per 900 bp (3/4 × 1/2 × (1/4)^3^ × 1/2 × 3/4 × 1/2) in the genome, it is possible that the promoter binding of HSF5 is tuned by cooperation with other DNA-binding factors (Fig. S5, Supplementary Data 5). Altogether, HSF5 acts as an atypical HSF transcription factor so that it executes a more pronounced role in regulating developmental genes rather than stress response genes through different mechanisms of DNA binding from canonical HSFs.

### HSF5 plays an essential role in pachytene progression during spermatogenesis

In males, meiotic prophase is accompanied by significant changes in gene expression programs (Schultz et al. 2003) (Shima et al. 2004) (Namekawa et al. 2006) (Green et al. 2018; Grive et al. 2019). At the pachytene stage in males, the progression of meiotic prophase is monitored under several layers of regulation, such as the pachytene checkpoint (Li et al. 2009) and meiotic sex chromosome inactivation (MSCI) (Burgoyne et al. 2009) (Ichijima et al. 2012) (Turner 2015). Concurrent with surveillance mechanisms, multiple gene regulatory programs are imposed on the male meiotic prophase to circumvent the barrier at the pachytene stage (Li et al. 2013) (Sin et al.2015) (da Cruz et al. 2016) (Ernst et al. 2019).

Our study revealed that HSF5 is a spermatocyte-specific transcription factor essential for progression beyond the pachytene stage. HSF5 binds to genes associated with biological processes of chromatin organization, such as *Kat6b, Kmt2d, Kmt3d, Kmt5b, Jmjd1c, Kdm3a, Atad2, Cbx7, Ep400, Ncor1, Rsbn1, Bend3,* which are generally expressed in a broad range of cell types (Fig 6). Our IP-MS analysis suggests that HSF5 interacts with KCTD19 (Fig S5), which has been shown to form a ZFP541-HDAC1/2-containing repressive complex (Horisawa-Takada et al. 2021), and we reasoned that HSF5 plays a role at least in repressing these target genes during pachytene progression. This is consistent with previous studies showing that pre-pachytene gene expression programs are suppressed by the ZFP541-KCTD19-containing repressive complex at the pachytene exit (Horisawa-Takada et al. 2021) (Oura et al. 2021) (Xu et al. 2022). In addition, HSF5 may positively regulate gene expression in cooperation with other transcription factors. Presumably, the absence of HSF5 indirectly delays DSB repair processes at the pachytene stage as a secondary effect of chromatin disorganization and aberrant gene expression (Fig 3). Therefore, HSF5 may trigger the reconstruction of the transcription network to promote pachytene progression and facilitate spermatid production. Our study sheds light on the regulatory mechanisms of gene expression that promote the developmental progression of the meiotic prophase, leading to spermatid differentiation.

## Acknowledgments

The authors thank Drs. Shosei Yoshida, Hitoshi Niwa for discussion, Yuki Takada for technical assistance, Akihiko Sakashita and Satoshi Namekawa for sharing their technical information of DCV FACS sorting, and Marry Ann Handel for provision of H1t antibody. This study was supported by the program of the Research for Inter-University Research Network for High Depth Omics, IMEG, Kumamoto University.

## Funding

This work was supported in part by KAKENHI grant (#20K22638) to R.S.; KAKENHI grant (#23K05641) to H.A.; KAKENHI grants (#19H05743, #20H03265, #20K21504, #22K19315, #23H00379, #16H06276 AdAMS, #22H04922 AdAMS) to K.I.; Grant from AMED PRIME (23gm6310021h0003) to K.I.; Grants from The Mitsubishi Foundation; The Naito Foundation; Astellas Foundation for Research on Metabolic Disorders; The Uehara Foundation; Takeda Science Foundation to K.I.

## Author contributions

S.Y., supervised by T.O. and E.K., performed most of experiments and wrote the draft of the manuscript.

R.S. performed scRNA-seq analysis, EMSA, informatics of RNA-seq and ChIP-seq data.

S.K. T.S. performed SEC-MALS analysis.

K.K. H.A. assisted cytological analysis.

S.I. assisted plasmid construction.

K.A. generated mutant mice and performed IVF.

N.T. performed MS analysis.

S.F. performed histological analysis.

K.Y. and S.U. assisted scRNA-seq and RNA-seq.

K.I. conducted the study and wrote the manuscript. The experimental design and interpretation of data were conducted by S.Y. R.S. and K.I.

## Declaration of interests

Authors declare no competing interests.

## Supplementary Figure legend

**Supplementary Figure 1. Specific expression of *Hsf5* orthologs in mouse and human testis. (related to Figure 1)**

**(A)** The tissue expression atlas of mouse *Hsf* gene paralogs are adapted from Expression Atlas (https://www.ebi.ac.uk/gxa/home). The expression levels (TPM : transcripts per million) are shown with the indicated color codes.

**(B)** UMAP plots show expression patterns of mouse *Hsf* gene paralogs and other key developmental genes are reanalyzed using public scRNA-seq data of spermatogenic cells in adult mouse testis (GEO: GSE109033) (Hermann et al. 2018). Key developmental marker genes include *Stra8*: differentiating spermatogonia and pleleptotene spermatocyte, *Meiosin*: pleleptotene spermatocyte, *Spo11*, *Terb1*: meiotic prophase spermatocyte, *Mybl1*: pachytene spermatocyte, *Prm1*: round and elongated spermatid.

**Supplementary Figure 2. Phenotypic analyses of *Hsf5* KO females. (related to Figure 2)**

**(A)** Hematoxylin and Eosin stained sections of WT, *Hsf5*+/- and *Hsf5* KO ovaries (8-weeks old). Biologically independent mice (N=3) for each genotype were examined. Scale bar: 500μm.

**(B)** Cumulative number of pups born from *Hsf5*+/- (n=3, all 4-weeks old at the start point of mating) and *Hsf5* KO (n=3, all 4-weeks old at the start point of mating) females.

**(C)** Fertility of *Hsf5* +/- and *Hsf5* KO females was examined by mating with WT males for the indicated period. Litter size is shown on the graph (Mean with SD).

**Supplementary Figure 3. Fluorescent sorting of meiotic prophase enriched spermatocytes by DCV fluorescence and light scattering for SMART RNA-seq (related to Figure 4)**

For SMART RNA-seq, meiotic prophase spermatocytes were isolated from **(A)** WT and **(B)** *Hsf5* KO testes at P17 by fluorescent sorting with DCV staining. **(A-1)(A-2)(B- 1)(B-2)** Debris and non-single cells were excluded by light scattering. **(A-3)(B-3)** Unstained cell and side population exclusion based on DCV fluorescence. **(A-4)(B-4)** DNA content determination based on DCV-blue fluorescence. **(A -5)(B -5)** Gating on E as a population with DNA content of 4C based on DCV-blue/DCV-red fluorescence. **(A -6)(B-6)** Back-gating of Gate E from the DCV plot on an FSC/BSC plot. We isolated cells gating on H as early meiotic prophase spermatocytes (Yeh et al. 2021). Precise gating of 4C testicular populations was confirmed by SYCP3+ positive immunostaining.

**Supplementary Figure 4. Clustering analysis of scRNA-seq data of WT and *Hsf5* KO spermatogenic germ cells (related to Figure 5)**

**(A)** Summary table of the 10X Genomics Chromium metrics for scRNA-seq analysis with WT and *Hsf5* KO testicular germ cells (P16). Indicated numbers of testes were pooled. Total number of testicular germ cells that were subjected to RNA-seq analysis and separated from other testicular somatic cells are shown. Percentages of the extracted testicular germ cells per total single cells that were subjected to RNA-seq analysis are shown.

**(B)** UMAP representation of scRNA-seq transcriptome profiles for testicular cells from P16 WT and *Hsf5* KO testes.

**(C)** Heat plot for the top 10 representative genes of each cluster. The relative expression levels are indicated with the color codes.

**(D)** Gene enrichment analysis of DEGs in the UMAP-defined cell clusters.

**Supplementary Figure 5. MS analyses of HSF5 interacting factors in testis extracts (related to Figure 6)**

**(A)** The immunoprecipitates (IP) by two different HSF5-N antibodies (HSF5-N1, HSF5-N2) from the chromatin-bound fractions of the testis extracts were subjected to liquid chromatography tandem-mass spectrometry (LC-MS/MS) analyses. The proteins identified by the LC-MS/MS analysis of HSF5N-IP are presented after excluding the proteins detected in the control IgG-IP. The proteins that were detected in both HSF5-N1 IP and HSF5-N2 IP with more than 1 different peptide hits are listed with SwissProt accession number, Amino acid length, the number of peptide hits and Mascot scores.

**(B)** The immunoprecipitates (IP) by HSF5-C antibody from the chromatin-bound fractions of the testis extracts. The proteins identified by the LC-MS/MS analysis of HSF5C-IP are presented as in A. See also Supplementary Data 5 for the raw data of LC-MSMS analysis.

Note that SMARCA5 and KCTD19 was repeatedly identified in the immunoprecipitates by HSF5-N1, HSF5-N2 HSF5-C antibodies.

**Supplementary Figure 6. Sequence alignment of DNA-binding domain of the mouse HSFs (related to Figure 6)**

**(A)** Amino acid sequences of DNA-binding domain in the mouse HSFs are aligned. HSF1, HSF2, and HSF4 possess a DNA recognition helix (red), containing a conserved Ser-Phe-Val-Arg-Gln amino acid sequence, which is known to insert into the major groove of the HSE. HSF5 possesses a Ser-Phe-Ile-Arg-Gln amino acid sequence at the corresponding position. HSF5 possesses insertion of amino acid sequence between the helix 2 and DNA recognition helix.

**(B)** Ribbon models of HSF5-DBD (1-167 a.a.) is shown that was predicted from AlphaFold2. Helixes are colored as shown in (A).

**Supplementary Figure 7. Uncropped images of gels and blots. (related to Figure 1, 2, 6)**

Full-length / uncropped images of agarose gel (Fig1D), western blot (Fig2B) and the scanned images of autoradiograph (Fig6F, Fig6G) are shown.

**Supplementary Data 1. Primers and oligos used in this study.**

**Supplementary Data 2. The list of DEGs in the SMART RNA-seq of the control versus *Hsf5* KO spermatocytes.**

Tab 1 : The complete list of expression profile of all genes that were detected by SMART RNA-seq.

Tab 2 : The complete result of up-regulated genes in *Hsf5* KO spermatocytes.

Tab 3 : The complete result of down-regulated genes in *Hsf5* KO spermatocytes.

Tab 4 : The complete result of Gene enrichment analysis for the up-regulated genes in *Hsf5* KO spermatocytes.

Tab 5 : The complete result of Gene enrichment analysis for the down-regulated genes in *Hsf5* KO spermatocytes.

Tab 6 : The down-regulated genes in *Hsf5* KO that are shown in the heatmap.

Tab 7-11 : The complete result of Gene enrichment analysis for the down-regulated genes in *Hsf5* KO spermatocytes in the clusters 1-5.

**Supplementary Data 3. The list of DEGs in the clusters of scRNA-seq of testicular germ cells.**

Shown are the complete list of DEGs in the clusters of scRNA-seq at P16. The complete result of Gene enrichment analysis for the highly expressed genes in the clusters are shown in other tabs (C0-C11).

**Supplementary Data 4. The gene list of HSF5-ChIP targets**

Tab 1 : The complete result of HSF5-ChIP targets.

Tab 2 : The complete result of Gene enrichment analysis for the HSF5-ChIP target genes.

Tab 3 : The HSF5-bound genes (Class1-7) that are shown in the heat map of the hierarchical clustering.

Tab 4 : The HSF1- and HSF2- bound genes in the testis are shown.

Tab 5 : The HSF1- and HSF2- bound gene sets that were described in the originally reported papers in previous ChIP on Chip studies (Akerfelt et al. 2008), (Akerfelt et al. 2010).

**Supplementary Data 5. The complete result of HSF5 IP-MS results**

Tab 1 : The complete list of HSF5-N1 and HSF5-N2 IP proteins from chromatin-bound fractions of the testis extracts identified by LC-MSMS.

Tab 2 : The complete list of HSF5-N1 and HSF5-N2 IP proteins identified from cytosol fractions of the testis extracts by LC-MSMS.

Tab 3 : The complete list of HSF5-C IP proteins identified from chromatin-bound fractions and cytosol fractions of the testis extracts by LC-MSMS.

Raw data are shown in other tabs

## Resource Availability

### Lead Contact

Further information and requests for the resources and reagents should be directed to and will be fulfilled by the Lead Contact, Kei-ichiro Ishiguro.

### Materials availability

Further information and requests for resources and reagents should be directed to and will be fulfilled by the Lead Contact, Kei-ichiro Ishiguro. Mouse lines generated in this study have been deposited to Center for Animal Resources and Development (CARD ID3259 for *Hsf5* knockout mouse #21 C57BL/6-*Hsf5^em1^*, CARD ID3268 for *Hsf5* knockout mouse #5 (C57BL/6-*Hsf5^em2^*), CARD ID3269 for *Hsf5* knockout mouse #9 (C57BL/6-*Hsf5^em3^*). Sequencing data have been deposited in DDBJ Sequence Read Archive (DRA) under the accession DRA 017033 for scRNA-seq data of P16 WT and *Hsf5* knockout testes and DRA017058 for the SMART RNA-seq data of the sorted meiotic prophase spermatocytes and DRA017059 for ChIP-seq data of WT testes. The antibodies are available upon request. There are restrictions to the availability of antibodies due to the lack of an external centralized repository for its distribution and our need to maintain the stock. We are glad to share antibodies with reasonable compensation by requestor for its processing and shipping. All unique/stable reagents generated in this study are available from the Lead Contact with a completed Materials Transfer Agreement.

## Data availability

All data supporting the conclusions are present in the paper and the supplementary materials. Raw sequence data generated in this study were publicly available as of the date of publication. Accession numbers are listed in the key resource table. Any additional information required to reanalyze the data reported in this paper is available from the lead contact upon request.

## Experimental Model and Subject Details

## Animals

*Hsf5* knockout (*Hsf5* KO) mice were with the C57BL/6 background. Male mice were used for ChIP-seq (age : postnatal day 10-20 old), immunoprecipitation of testis extracts (age : postnatal day 18-23 old, 4-weeks and 8-weeks old), histological analysis of testes, and immunostaining of testes, RNA extraction (age : postnatal day 15-21, 4-weeks and 8-weeks old). Female mice were used for histological analysis of the ovaries and RNA extraction (age : postnatal 4-weeks and 8-weeks old) (embryonic day 12.5-18.5). Whenever possible, each knockout animal was compared among littermates or age-matched non-littermates from the same colony, unless otherwise described. Animal experiments were approved by the Institutional Animal Care and Use Committee (approval F28-078, A2022-001).

## Method Details

### Generation of *Hsf5* knockout mice and genotyping

*Hsf5* knockout mice were generated by introducing Cas9 protein (317-08441; NIPPON GENE, Toyama, Japan), tracrRNA (GE-002; FASMAC, Kanagawa, Japan), synthetic crRNA (FASMAC), and ssODN into C57BL/6N fertilized eggs using electroporation. For the generation of *Hsf5* Exon1-6 deletion (Ex1-6Δ) allele, the synthetic crRNAs were designed to direct CCTTAAATTCAAATTAGATG(AGG) of the 5’upstream of *Hsf5* exon 1 and ATGTAGACAAAAGCACTGAG(AGG) in the exon 6. ssODN: 5’-AAAAATCTAAAATAAGAAAACAGTGTTAACCTCTCATGAGGTCCTTGGTACCTGGCAGAAGGGAATAAAG -3’ was used as a homologous recombination template.

The electroporation solutions contained tracrRNA(10μM), synthetic crRNA(10μM), Cas9 protein (0.1 μg/μl), ssODN (1μg/μl) for *Hsf5* knockout in Opti-MEM I Reduced Serum Medium (31985062; Thermo Fisher Scientific). Electroporation was carried out using the Super Electroporator NEPA 21 (NEPA GENE, Chiba, Japan) on Glass Microslides with round wire electrodes, 1.0 mm gap (45-0104; BTX, Holliston, MA). Four steps of square pulses were applied (1, three times of 3 mS poring pulses with 97 mS intervals at 30 V; 2, three times of 3 mS polarity-changed poring pulses with 97 mS intervals at 30 V; 3, five times of 50 mS transfer pulses with 50 mS intervals at 4 V with 40% decay of voltage per each pulse; 4, five times of 50 mS polarity-changed transfer pulses with 50 mS intervals at 4 V with 40% decay of voltage per each pulse). The targeted *Hsf5* Exon1-6Δ allele in F0 mice were identified by PCR using the following primers; Hsf5-g-F1: 5’-GGGAGATCATAGCTGGTCATTAAGC-3’ and Hsf5-g-R2: 5’-CAGAGGGATAAGAAAATTGGTGATAG-3’ for the knockout allele (300bp). Hsf5-g-F1 and Hsf5-g-R1: 5’-TCTCCCACCGTTCTCGATCC-3’ for the Ex1 of WT allele (678 bp). The PCR amplicons were verified by Sanger sequencing. Primer sequences are listed in Supplementary Data 1.

### Histological Analysis

For hematoxylin and eosin staining, testes, epididymis and ovaries were fixed in 10% formalin or Bouin solution, and embedded in paraffin. Sections were prepared on CREST-coated slides (Matsunami) at 6 μm thickness. The slides were dehydrated and stained with hematoxylin and eosin.

For Immunofluorescence staining, testes were embedded in Tissue-Tek O.C.T. compound (Sakura Finetek) and frozen. Cryosections were prepared on the CREST-coated slides (Matsunami) at 8 μm thickness, and then air-dried and fixed in 4% paraformaldehyde in PBS at pH 7.4. The serial sections of frozen testes were fixed in 4% PFA for 5 min at room temperature and permeabilized in 0.1% TritonX100 in PBS for 10 min. The sections were blocked in 3% BSA/PBS or Blocking One (Nakarai), and incubated at room temperature with the primary antibodies in a blocking solution. After three washes in PBS, the sections were incubated for 1 h at room temperature with Alexa-dye-conjugated secondary antibodies (1:1000; Invitrogen) in a blocking solution. TUNEL assay was performed using MEBSTAIN Apoptosis TUNEL Kit Direct (MBL 8445). DNA was counterstained with Vectashield mounting medium containing DAPI (Vector Laboratory). Statistical analyses, and production of graphs and plots were done using GraphPad Prism9 or Microsoft Excel.

### Imaging

Immunostaining images were captured with DeltaVision (Cytiva). The projection of the images was processed with the SoftWorx software program (Cytiva). All images shown were Z-stacked. Excitation intensity and exposure time were adjusted for each condition, and images were simultaneously acquired at the same condition for comparable analyses. The brightness of images was linearly adjusted to the same range of scale on the signal intensity histogram using SoftWorx software program for better visibility. For counting seminiferous tubules, immunostaining images were captured with BIOREVO BZ-X710 (KEYENCE), and processed with BZ-H3A program. XY-stitching capture by 10x objective lens was performed for multiple-point color images using BZ-X Wide Image Viewer. Images were merged over the field using BZ-H3A Analyzer (KEYENCE). Bright field images were captured with OLYMPUS BX53 fluorescence microscope and processed with CellSens standard program.

### Production of antibodies against HSF5

Polyclonal antibodies against mouse HSF5 N-terminal (aa1-209) and HSF5 C-terminal (aa324-624) were generated by immunizing rabbits. His-tagged recombinant proteins of HSF5 N-terminal region (aa1-209) and HSF5 C-terminal region (aa324-624) were produced by inserting cDNA fragments in-frame with pET19b (Novagen) respectively in *E. coli* strain BL21-CodonPlus (DE3)-RIPL (Agilent), solubilized in a denaturing buffer (6 M HCl-Guanidine, 20 mM Tris-HCl pH 7.5) and purified by Ni-NTA (QIAGEN) under denaturing conditions. The antibodies were affinity-purified from the immunized serum with immobilized antigen peptides on CNBr-activated Sepharose (GE healthcare).

### Antibodies

The following antibodies were used for immunoblot (IB) and immunofluorescence (IF) studies: Gunia pig anti-SYCP3 (IF, 1:2000, our home made) (Ishiguro et al. 2020), Rat anti-SYCP3 (IF, 1:1000, our home made) (Ishiguro et al. 2020), Rabbit anti-SYCP1 (IF, 1:1000, Abcam ab15090), Mouse anti-γH2AX (IF, 1:1000, Abcam ab26350), Guinea pig anti-H1t (IF, 1:2000, kindly provided by Marry Ann Handel)(Cobb et al. 1999), Rabbit α-tubulin DM1A (IB, 1:2000, Sigma 05-829), rabbit anti-HSF5 N (IB,IF, 1:1000, our home made)(this paper), rabbit anti-HSF5 C (IB,IF, 1:1000, our home made)(this paper), Mouse anti-FLAG M2 (IB, 1:1000, Sigma-Aldrich F1804), Rabbit anti-HA (IB,IF, 1:1000, Abcam ab9110), Mouse Anti-MLH1 (IF, 1:200, Cell Signaling 3515), Rabbit anti-DMC1 (IF, 1:500, Santa Cruz: SC-22768), Rabbit BRCA1 (IF, 1:2000, kindly provided by Satoshi Namekawa) (Tanno et al. 2022).

Following secondary antibodies were used : Goat anti-Rabbit IgG Alexa 488 (IF, 1:1000, Invitrogen 31570), Goat anti-Gunia pig IgG-Alexa Fluour 488 (IF, 1:1000, Abcam ab150185), Donkey Anti-Mouse IgG Alexa 555 (IF, 1:1000, Invitrogen A31570), Goat Anti-Rat IgG Alexa 555 (IF, 1:1000, Thermo Fisher A21434), Goat anti-Gunia pig IgG-Alexa Fluour 555 (IF, 1:1000, Abcam ab150186), Goat anti-rabbit IgG-Alexa Fluour 647 (IF, 1:1000, Thermo Fisher, A21244), Donkey anti-mouse IgG-Alexa Fluour 647 (IF, 1:1000, Thermo Fisher, A31571), Sheep anti-mouse IgG-Horseradish Peroxidase (IB: 1/10000, Cytiva, NA931), Donkey anti-rabbit IgG-Horseradish Peroxidase (IB: 1/10000, Cytiva, NA934).

### PCR with reverse transcription

Total RNA was isolated from tissues and embryonic gonads using TRIzol (Thermo Fisher). cDNA was generated from total RNA using Superscript III (Thermo Fisher) followed by PCR amplification using Ex-Taq polymerase (Takara) and template cDNA. For RT-qPCR, total RNA was isolated from WT embryonic ovaries (E12.5 – E18.5, n = 3 for each), and WT testes (n = 3), and cDNA was generated as described previously (Horisawa-Takada et al. 2021). *Hsf5* cDNA was quantified by DCT method using TB Green Premix Ex Taq II (Tli RNaseH Plus) and Thermal cycler Dice (Takara), and normalized by *GAPDH* expression level.

Sequences of primers used for RT-PCR and RT-qPCR were as follows:

Hsf1_RTPCR_F1: 5’-GCCCCTCTTCCTTTCTGCAT-3’

Hsf1_RTPCR_R1: 5’-TCATGTCGGGCATGGTCAC-3’

Hsf2_RTPCR_F1: 5’-TCCTGTTAGCAGAAACGGCA-3’

Hsf2_RTPCR_R1: 5’-GGGATCACTTCCAAAGACGA-3’

Hsf3_RTPCR_F1: 5’-AGCTTGATCTCAGTGGGGGA-3’

Hsf3_RTPCR_R1: 5’-ACTAGCCAGCAGCCATTGAA-3’

Hsf4_RTPCR_F1: 5’-GAACTCAGGCAGCAGAACGA-3’

Hsf4_RTPCR_R1: 5’-GGAGGGGCGACTGGATAAAG-3’

Hsfy2_RTset_1F: 5’-AATGCAGGCTGTTTCCCCTA-3’

Hsfy2_RTset_1R: 5’-GGTATGCGGTGGCCTCTTTA-3’

Gapdh_F2(Gaphdh5599): 5’-ACCACAGTCCATGCCATCAC-3’

Gapdh_R2(Gapdh5600): 5’-TCCACCACCCTGTTGCTGTA-3’

HSF5_RTset1_F: 5’-AGGGCTACCATTCAGCACAC-3’

HSF5_RTset1_R: 5’-GACTTGTTAGCAGGCCCCAT-3’

Primer sequences are listed in Supplementary Data 1.

### Electrophoretic mobility shift assay (EMSA)

HSF5 N-terminal protein (aa1-209) was purified by Ni-NTA under native condition (20 mM Tris-HCl pH 7.5, 150mM NaCl, 300mM imidazole, 1% TritonX100). Purified protein was subjected to HiTrap Q column using AKTA pure system, and then the flowthrough fraction was collected. The flowthrough fraction was subjected to HiTrap S column, and eluted by salt gradient from 0 to 1 M NaCl in a buffer (20 mM Tris-HCl pH 8.0, 1mM 2-Mercaptoethanol). The eluted protein was dialyzed against a buffer (20 mM Tris-HCl pH 8.0, 100mM NaCl, 1mM 2-Mercaptoethanol, 10% glycerol). The annealed synthetic oligonucleotide DNA was labeled with [α-^32^P]dCTP (Perkin Elmer NEG013H250UC, 3000Ci/mmol) by Klenow polymerase. ^32^P-labeled DNA was separated by electrophoresis in a 10 % polyacrylamide gel, eluted from gel slices with an elution buffer containing 1 mM EDTA, and 10 mM Tris-HCl (pH 8.0), and precipitated by ethanol. The synthetic oligonucleotide DNA sequences are as followings; For the target (T) sequence probe, HSF5-EMSA-T-up : 5’-agggcatagatatggaactctcttagaactctcttagaacttactactg-3’ HSF5-EMSA-T-bottom : 5’-aggcagtagtaagttctaagagagttctaagagagttccatatctatgc-3’ For the mutant (M) sequence probe, HSF5-EMSA-M-up : 5’-agggcatagataaccttgagtctatcttgagtctatcttgaaactactg-3’. HSF5-EMSA-M-bottom : 5’-aggcagtagtttcaagatagactcaagatagactcaaggttatctatgc-3’ The ^32^P-labeled DNA (0.04 pmol, 2 × 10^4^ to 4 × 10^4^ cpm) was mixed with 18.75-300 ng of HSF5 DNA binding domain in 10 μl of a buffer containing 20mM Tris-HCl pH 8.0, 100mM NaCl and 2mM MgCl_2_. The mixture was incubated at 25°C for 10 min and loaded with glycerol dye mix (25% glycerol, 1 mM EDTA, 0.01% bromophenol blue) on an 8 % polyacrylamide gel (acrylamide bis-acrylamide, 29:1) containing 89 mM Tris-borate (pH 8.3)-2 mM EDTA. After electrophoresis, the gel was dried and subjected to autoradiography by the Typhoon FLA7000 Biomolecular imager (Cytiva).

### SEC–MALS Experiments

The full length HSF5 expression construct was cloned into a pMAL-c6T vector (New England Biolabs) and fused to His^6^-MBP and a tobacco etch virus protease cleavage site at the N-terminus. His-MBP-full length HSF5 was purified by HisTrap HP (Cytiva), and gel filtration using a Superdex 200 pg 16/600 column (Cytiva). Size exclusion chromatography with multi-angle light scattering (SEC–MALS) was performed using a DAWN HELEOS8+ (Wyatt Technology Corporation, Santa Barbara, CA, USA), a high-performance liquid chromatography pump LC-20AD (Shimadzu, Kyoto, Japan), a refractive index detector RID-20A (Shimadzu), and a UV–vis detector SPD-20A (Shimadzu), which were located downstream of the Shimadzu liquid chromatography system connected to a PROTEIN KW-803 gel filtration column (Cat. no. F6989103; Shodex, Tokyo, Japan). Differential RI (Shimadzu) downstream of MALS was used to determine the protein concentrations. The running buffer used contained 25 mM HEPES/KOH (pH 7.2) and 150 mM KCl. Approximately 100 μL of the sample was injected at a flow rate of 1.0 mL /min. Data was then analyzed using ASTRA version 7.0.1 (Wyatt Technology Corporation). Molar mass analysis was also performed over half of the width of the UV peak top height. 30 min after incubation at 42 °C, 100 μL of MBP-full length HSF5 (12.6 μM) or HSF5-DBD (26.6 μM) was injected.

### Preparation of testis extracts and immunoprecipitation

Testis chromatin-bound and -unbound extracts were prepared as described previously (Tani et al. 2022). Briefly, testes were removed from male C57BL/6 mice (P18-23, 4-weeks and 8-weeks old), detunicated, and then resuspended in low salt extraction buffer (20 mM Tris-HCl pH 7.5, 100 mM KCl, 0.4 mM EDTA, 0.1% TritonX100, 10% glycerol, 1 mM β-mercaptoethanol) supplemented with Complete Protease Inhibitor (Roche). After homogenization, the soluble chromatin-unbound fraction was separated after centrifugation at 100,000*g* for 10 min at 4°C. The chromatin bound fraction was extracted from the insoluble pellet by high salt extraction buffer (20 mM HEPES-KOH pH 7.0, 400 mM KCl, 5 mM MgCl_2_, 0.1% Tween20, 10% glycerol, 1 mM β-mercaptoethanol) supplemented with Complete Protease Inhibitor. The solubilized chromatin fraction was collected after centrifugation at 100,000*g* for 10 min at 4°C.

For immunoprecipitation of endogenous HSF5 from extracts, 5 µg of affinity-purified rabbit anti-HSF5-C, HSF5-N1, HSF5-N2 and control IgG antibodies were crosslinked to 50 µl of protein A-Dynabeads (Thermo-Fisher) by DMP (Sigma). The antibody-crosslinked beads were added to the testis extracts prepared above. The beads were washed with low salt extraction buffer. The bead-bound proteins were eluted with 40 µl of elution buffer (100 mM Glycine-HCl pH 2.5, 150 mM NaCl), and then neutralized with 4 µl of 1 M Tris-HCl pH 8.0. The immunoprecipitated proteins were run on 4-12 % NuPAGE (Thermo-Fisher) in MOPS-SDS buffer and immunoblotted. Immunoblot images were developed using ECL prime (GE healthcare) and captured by FUSION Solo (VILBER).

### Mass spectrometry

The immunoprecipitated proteins were run on 4-12 % NuPAGE (Thermo Fisher) by 1 cm from the well and stained with SimplyBlue (Thermo Fisher) for in-gel digestion. The gel containing proteins was excised, cut into approximately 1mm sized pieces. Proteins in the gel pieces were reduced with DTT (Thermo Fisher), alkylated with iodoacetamide (Thermo Fisher), and digested with trypsin and Lysyl endopeptidase (Promega) in a buffer containing 40 mM ammonium bicarbonate, pH 8.0, overnight at 37°C. The resultant peptides were analyzed on an Advance UHPLC system (ABRME1ichrom Bioscience) connected to a Q Exactive mass spectrometer (Thermo Fisher) processing the raw mass spectrum using Xcalibur (Thermo Fisher Scientific). The raw LC-MS/MS data was analyzed against the SwissProt protein/translated nucleotide database restricted to *Mus musculus* using Proteome Discoverer version 1.4 (Thermo Fisher) with the Mascot search engine version 2.5 (Matrix Science). A decoy database comprised of either randomized or reversed sequences in the target database was used for false discovery rate (FDR) estimation, and Percolator algorithm was used to evaluate false positives. Search results were filtered against 1% global FDR for high confidence level. All full lists of LC-MS/MS data are shown in Supplementary Supplementary Data 5 (Excel file).

### ChIP-seq Analysis

The seminiferous tubules from male C57BL/6 mice (P10-20) were minced and digested with accutase and 0.5 units/ml DNase II, followed by filtration through a 40 μm cell strainer (FALCON). Testicular cells were fixed in 1% formaldehyde (Thermo-Fisher) and 2mM Disuccinimidyl glutarate (ProteoChem) in PBS for 10 minutes at room temperature. Crosslinked cells were lysed with LB1 (50 mM HEPES pH 7.5, 140 mM NaCl, 1 mM EDTA, 10% glycerol, 0.5% NP-40, and 0.25% Triton X-100) and washed with LB2 (10 mM Tris-HCl pH 8.0, 200 mM NaCl, 1 mM EDTA, 0.5 mM EGTA). Chromatin lysates were prepared in LB3 (50 mM Tris-HCl pH8.0, 1% SDS, 10 mM EDTA, proteinase inhibitor cocktail (Sigma)), by sonication with Covaris S220 (Peak Incident Power, 175; Acoustic Duty Factor, 10%; Cycle Per Burst, 200; Treatment time, 600sec; Cycle, 6).

HSF5 ChIP was performed using chromatin lysates and protein A Dyna-beads (Thermo-Fisher) coupled with 5 μg of rabbit anti-HSF5-N and rabbit anti-HSF5-C antibodies, and rabbit control IgG as described previously (Horisawa-Takada et al. 2021). After 4 hours of incubation at 4 °C, beads were washed 4 times in a low salt buffer (20 mM Tris-HCl (pH 8.0), 0.1% SDS, 1% (w/v) TritonX-100, 2 mM EDTA, 150 mM NaCl), and two times with a high salt buffer (20 mM Tris-HCl (pH 8.0), 0.1% SDS, 1% (w/v) TritonX-100, 2 mM EDTA, 500 mM NaCl). Chromatin complexes were eluted from the beads by agitation in elution buffer (10 mM Tris-HCl (pH 8.0), 300 mM NaCl, 5 mM EDTA, 1% SDS) and incubated overnight at 65 °C for reverse-crosslinking. Eluates were treated with RNase A and Proteinase K, and DNA was ethanol precipitated. ChIP-seq libraries were prepared using 20 ng of input DNA, and 4 ng of ChIP DNA with KAPA Library Preparation Kit (KAPA Biosystems) and NimbleGen SeqCap Adaptor Kit A or B (Roche) and sequenced by Illumina Hiseq 1500 to obtain single end 50 nt reads.

ChIP-seq reads were trimmed to remove adapter sequence when converting to a fastq file. The trimmed ChIP-seq reads were mapped to the UCSC mm10 genome assemblies using Bowtie2 v2.3.4.1 with default parameters. Peak calling was performed using MACS program v2.1.0 (Zhang et al. 2008) (https://github.com/macs3-project/MACS) with the option (-g mm -p 0.00001). To calculate the number of overlapping peaks between HSF5-N ChIP and HSF5-C ChIP, we used bedtools program (v2.27.1)(Quinlan and Hall 2010) (https://bedtools.readthedocs.io/en/latest/). ChIP binding regions were annotated with BED file using Cis-regulatory Element System (CEAS) v0.9.9.7 (package version 1.0.2), in which the gene annotation table was derived from UCSC mm10. Motif identification was performed using MEME-ChIP v5.1.1 website (http://meme-suite.org/tools/meme-chip)(Bailey 2011). The motif database chosen was “JASPAR Vertebrates and UniPROBE Mouse”. BigWig files, which indicate occupancy of HSF5, were generated using deepTools (v3.1.0) and visualized with Integrative Genomics Viewer software (v.2.8.3) http://software.broadinstitute.org/software/igv/home. GO-term analyses were performed using DAVID Bioinformatics Resources 6.8 (Huang da et al. 2009) (https://david.ncifcrf.gov/). Aggregation plots and heatmaps for each sample against the dataset of HSF5 target genes were made using deeptools program v3.5.0. The nearest genes of the ChIP-seq peaks were determined by GREAT website (v4.0.4) (http://great.stanford.edu/public/html/). Sequencing data are available at DDBJ Sequence Read Archive under the accession DRA017059.

### SMART RNA-seq Analysis

Spermatocytes at zygotene/pachytene from WT and *Hsf5* KO male testis were sorted using SH800S cell sorter (SONY) using DyeCycle Violet (DCV) stain (Thermo Fisher)(Yeh et al. 2021). Total RNAs were prepared by Trizol (Thermo Fisher) and quality of total RNA was confirmed by BioAnalyzer 2100 (RIN > 8) (Agilent). cDNA was synthesized by SMART-Seq mRNA kit (Takara, 634772). Library DNAs were prepared according to the Nextera XT DNA Library Preparation Kit (Illumina, FC-131-1096) and sequenced by Illumina NextSeq 500 (Illumina) using Nextseq 500/550 High Output v2.5 Kit (Illumina) to obtain single end 75 nt reads.

Resulting reads were aligned to the mouse genome UCSC mm10 using STAR ver.2.7.8a after trimmed to remove adapter sequence and low-quality ends using Trim Galore! v0.6.6 (cutadapt v2.5). Differential expression analysis using TPM was done by RSEM v1.3.3, and DESeq2 (v1.36.0). GTF file was derived from UCSC mm10. Principal component analysis and GO-term analysis were performed using DAVID Bioinformatics Resources 6.8 (Huang da et al. 2009). Sequencing data are available at DDBJ Sequence Read Archive under the accession DRA017058.

### Single-cell RNA-sequencing

Testes were collected from WT and *Hsf5* KO at P16. Testes were minced in 1.2mg/ml type I collagenase (Wako) and incubated for 10min. at 37°C, then tissue pellets were collected by centrifugation. Accutase (Innovative Cell Technologies, Inc.) was added to tissue pellet and incubated for 5min. at 37°C. After incubation, DMEM with 10% FBS was added to block Accutase and aggregate was disrupted by pipetting. Then, cell suspensions were filtered through a 95-μm and 35-μm cell strainer sieve (BD Bioscience). Cells were collected by centrifugation and re-suspended in PBS containing 0.1% BSA, and cells were collected using Cell Sorter SH800 (SONY) to remove dead cells. Collected cells were re-suspended in DMEM containing 10% FBS. Resulting approximate 10,000 single-cell suspensions were loaded on Chromium Controller (10X Genomics Inc.). Single cell RNA-seq libraries were generated using Chromium Single Cell 3’ Reagent Kits v3 following manufacturer’s instructions, and sequenced on an Illumina HiSeq X to acquire paired end 150 nt reads. The number of used embryos, the total numbers of single cells captured from testes, mean depth of reads per cell, average sequencing saturation (%), the number of detected genes, median UMI counts/cell, total number of cells before and after QC, are shown in Figure S5. Sequencing data are available at DDBJ Sequence Read Archive (DRA) under the accession DRA 017033.

### Statistical analysis of scRNA-seq

Fastq files were processed and aligned to the mouse mm10 transcriptome (GENCODE vM23/Ensembl 98) using the 10X Genomics Cell Ranger v 4.0 pipeline. Further analyses were conducted on R (ver.3.6.2) via RStudio (ver.1.2.1335). Quality assessment of scRNA-seq data and primary analyses were conducted using the Seurat package for R (v3.2.2) (Butler et al. 2018) (Stuart et al. 2019). Only the cells that expressed more than 200 genes were used for further analysis to remove the effect of low-quality cells. The scRNA-seq data were merged and normalized using SCTransform function built in Seurat. The dimensional reduction analysis and visualization of cluster were conducted using RunUMAP function built in Seurat. The clustering of cells was conducted using FindNeighbors and FindClusters built in Seurat with default setting. Determination of differentially expressed genes (DEGs) was performed using FindAllMarkers function built in Seurat with following options, only.pos = T, min.pict = 0.25 and logfc.threshold = 0.25, for the identification of marker genes in each cluster, and FindMarker function built in Seurat with default settings to characterize the arbitrary group pf cells. To extract the germ cell population, the cell populations expressing germ cell marker, *Ddx4*, were selected. The selected cell populations were re-plotted on the UMAP. The cell populations showing the expression of somatic cell marker genes and the enrichment of mitochondrial genes were removed and the remaining cells were used as germ cell population. RNA velocity analysis was conducted using the RNA velocyto package for python (Python 3.7.3) and R with default settings (La Manno et al. 2018), and visualized on UMAP plots built in Seurat.

### Gene set enrichment analysis

Gene set enrichment analyses were performed using Metascape (Zhou et al. 2019) with default settings and the results were visualized using R and GraphPad Prism9. To characterize the feature of each cluster, top100 representative genes in each cluster were used.

### Public ChIP-seq Data and RNA-seq data Analysis

MEIOSIN ChIP-seq data described in our previous study (Ishiguro et al. 2020) was analyzed for the *Hsf5* locus. MEIOSIN binding site was shown along with genomic loci from Ensembl on the genome browser IGV. 10xGenomics scRNA-seq data of of mouse adult testis was derived from GEO: GSE109033 (Hermann et al. 2018). Reanalyses of scRNA-seq data were conducted using the Seurat package for R (v.3.1.3) (Stuart et al. 2019) and pseudotime analayses were conducted using monocle package for R: R (ver. 3.6.2), RStudio (ver.1.2.1335), and monocle (ver. 2.14.0) (Qiu et al. 2017) following developer’s tutorial. The tissue expression atlas of mouse *Hsf* gene paralogs are adapted from Expression Atlas (https://www.ebi.ac.uk/gxa/home), and the expression levels are shown using Microsoft Excel.

### Quantification and Statistical analysis

Statistical analyses, and production of graphs and plots were done using GraphPad Prism9 or Microsoft Excel (version 16.48).

## Notes

### Competing Interest Statement

The authors have declared no competing interest.

