## Supplemental figures for "Atypical heat shock transcription factor HSF5 is critical for male meiotic prophase under non-stress conditions"

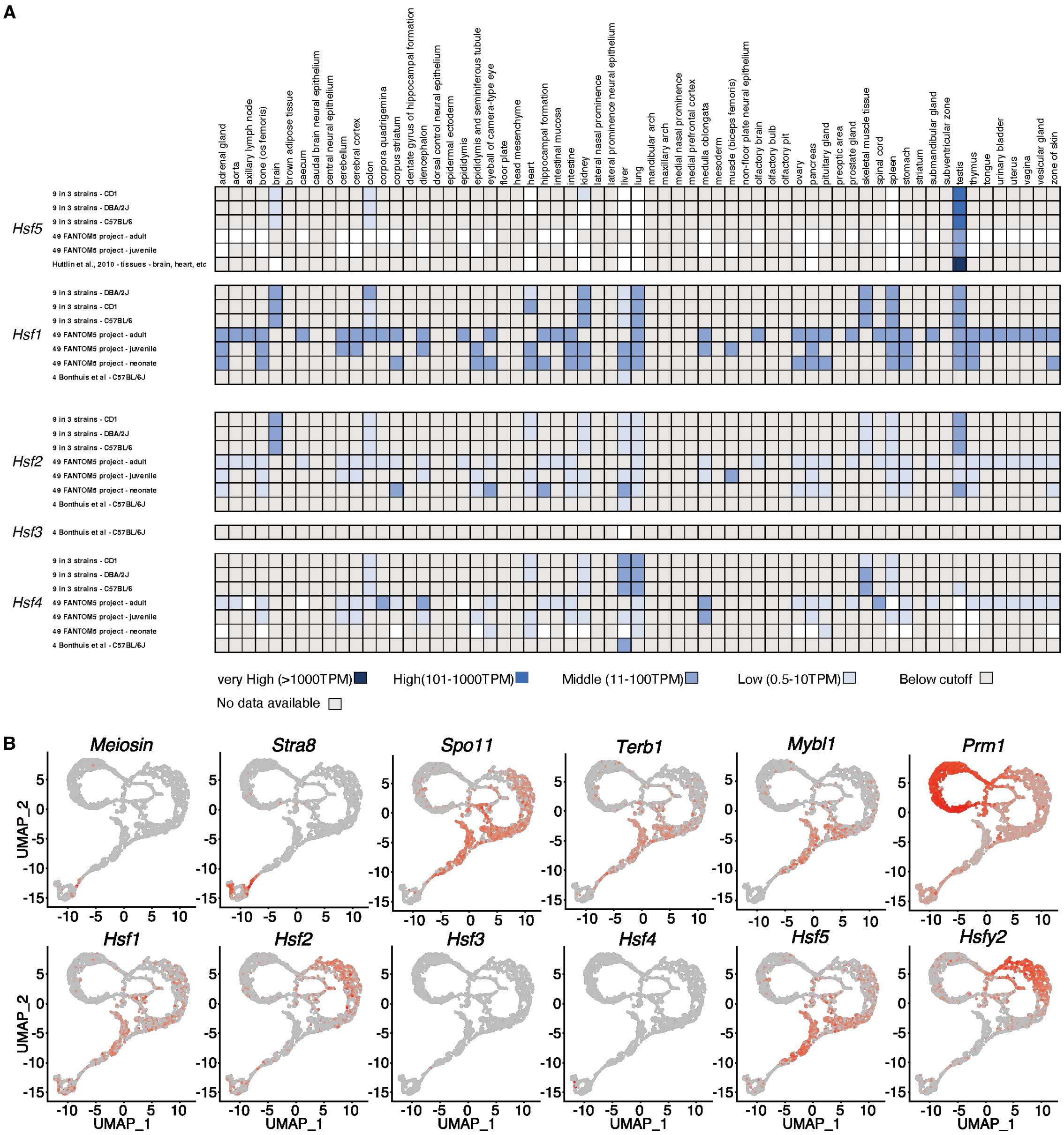


**Supplementary Figure 1. Specific expression of *Hsf5* orthologs in mouse and human testis. (related to Figure 1)**

**(A)** The tissue expression atlas of mouse *Hsf* gene paralogs are adapted from Expression Atlas (https://www.ebi.ac.uk/gxa/home). The expression levels (TPM : transcripts per million) are shown with the indicated color codes.

**(B)** UMAP plots show expression patterns of mouse *Hsf* gene paralogs and other key developmental genes are reanalyzed using public scRNA-seq data of spermatogenic cells in adult mouse testis (GEO: GSE109033) (Hermann et al. 2018). Key developmental marker genes include *Stra8*: differentiating spermatogonia and pleleptotene spermatocyte, *Meiosin*: pleleptotene spermatocyte, *Spo11*, *Terb1*: meiotic prophase spermatocyte, *Mybl1*: pachytene spermatocyte, *Prm1*: round and elongated spermatid.

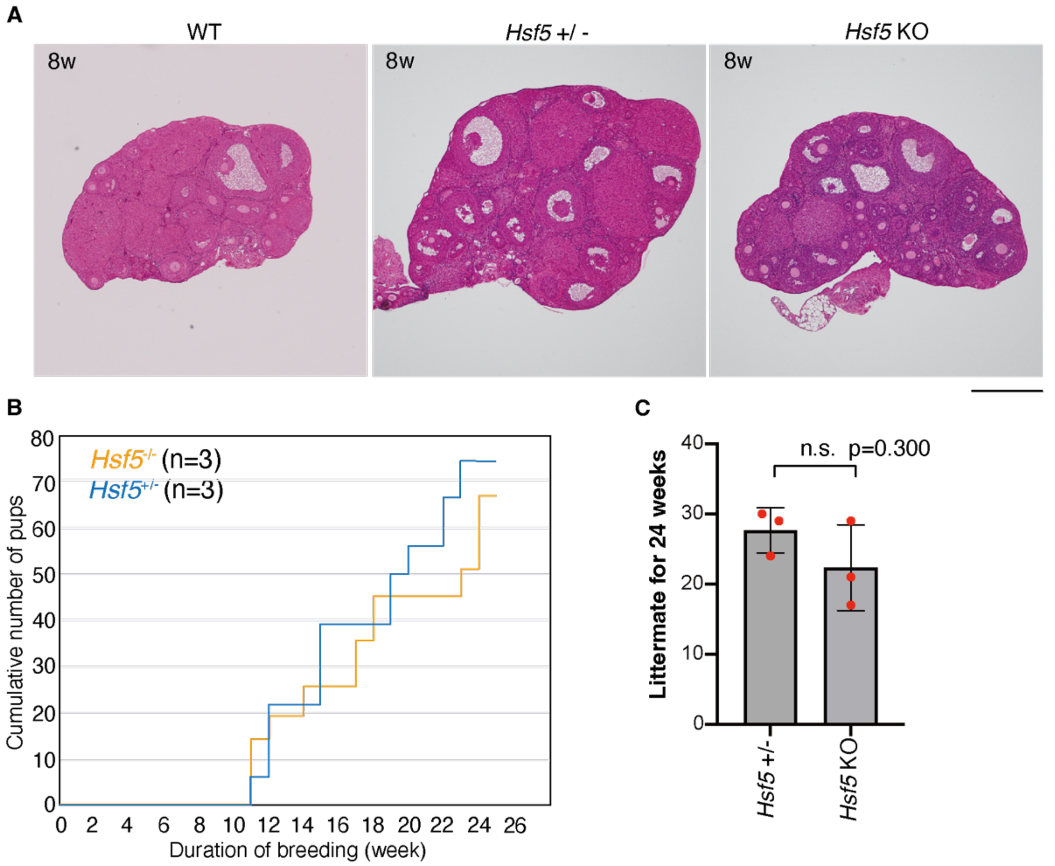


**Supplementary Figure 2. Phenotypic analyses of *Hsf5* KO females. (related to Figure 2)**

**(A)** Hematoxylin and Eosin stained sections of WT, *Hsf5*+/- and *Hsf5* KO ovaries (8-weeks old). Biologically independent mice (N=3) for each genotype were examined. Scale bar: 500μm.

**(B)** Cumulative number of pups born from *Hsf5*+/- (n=3, all 4-weeks old at the start point of mating) and *Hsf5* KO (n=3, all 4-weeks old at the start point of mating) females.

**(C)** Fertility of *Hsf5* +/- and *Hsf5* KO females was examined by mating with WT males for the indicated period. Litter size is shown on the graph (Mean with SD).

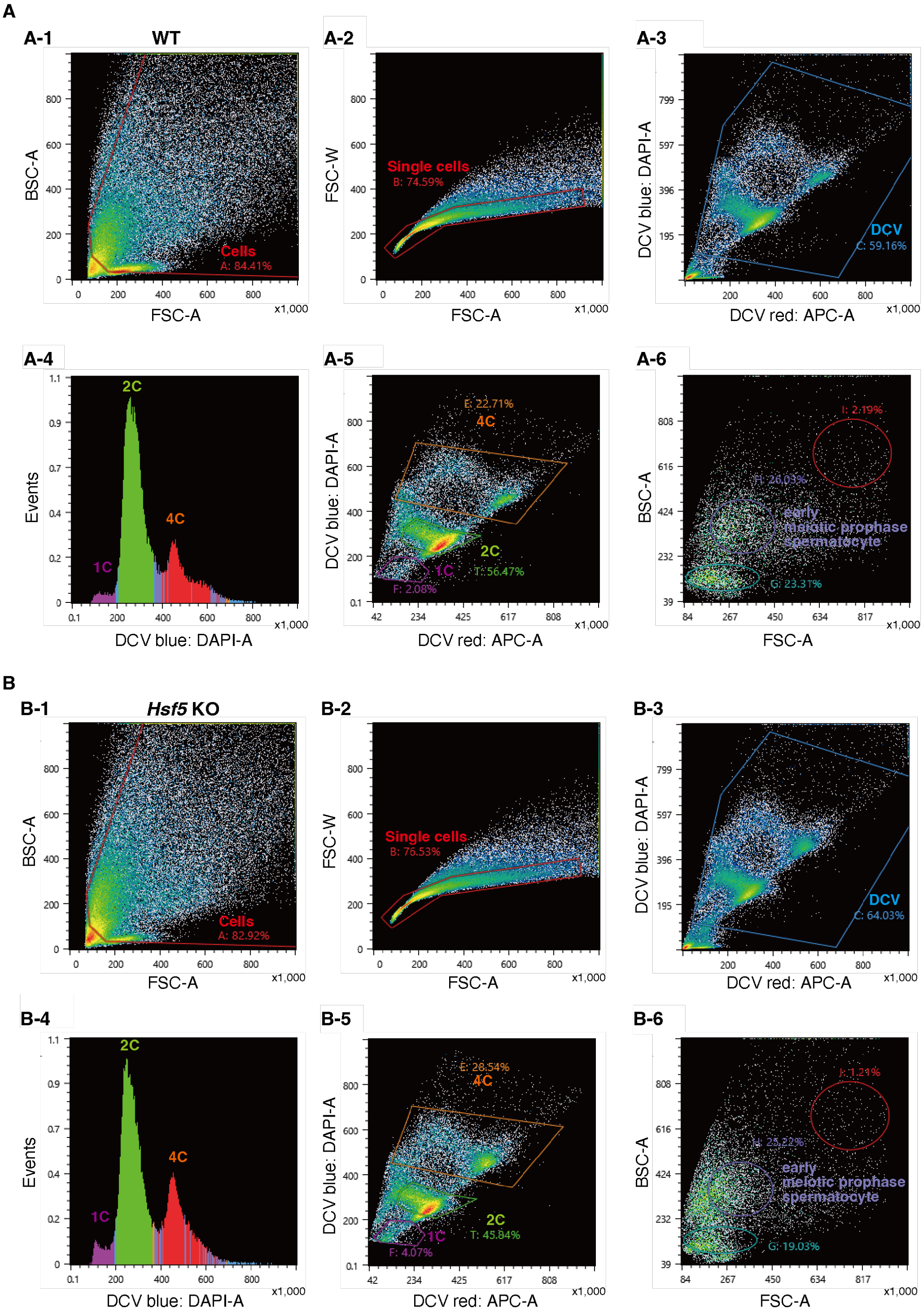


**Supplementary Figure 3. Fluorescent sorting of meiotic prophase enriched spermatocytes by DCV fluorescence and light scattering for SMART RNA-seq (related to Figure 4)**

For SMART RNA-seq, meiotic prophase spermatocytes were isolated from **(A)** WT and **(B)** *Hsf5* KO testes at P17 by fluorescent sorting with DCV staining. **(A-1)(A-2)(B-1)(B-2)** Debris and non-single cells were excluded by light scattering. **(A-3)(B-3)**

Unstained cell and side population exclusion based on DCV fluorescence. **(A-4)(B-4)** DNA content determination based on DCV-blue fluorescence. **(A-5)(B-5)** Gating on E as a population with DNA content of 4C based on DCV-blue/DCV-red fluorescence. **(A-6)(B-6)** Back-gating of Gate E from the DCV plot on an FSC/BSC plot. We isolated cells gating on H as early meiotic prophase spermatocytes (Yeh et al. 2021). Precise gating of 4C testicular populations was confirmed by SYCP3+ positive immunostaining.

**A**

| | Platform: Chromium (10x Genomics) | |
|---|---|---|
| Sampling stage: | P16 | |
| Genotype: | wild type | *Hsf5*-KO |
| Cell type: | whole testicular cells | |
| Number of used indivisuals: | 3 | 3 |
| **Total number of genes detected:** | 22,518 | 22,106 |
| Sequencing Saturation: | 72.6% | 70.3% |
| Mean reads / cell: | 40,968 | 32,472 |
| Median genes / cell: | 1,312 | 1,278 |
| Median UMI counts / cell: | 2,918 | 2,673 |
| **Total number of cells:** | 9,275 | 11,023 |
| Number of analyzed testicular germ cells: | 1,087 (11.7%) | 1,911 (17.3%) |

**B**

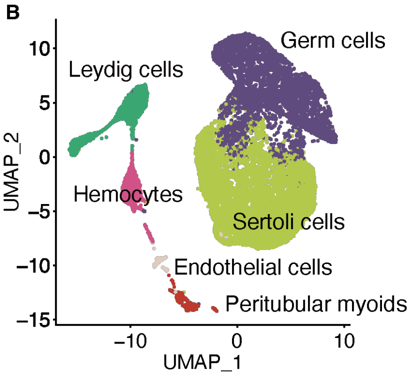


**C** **D**

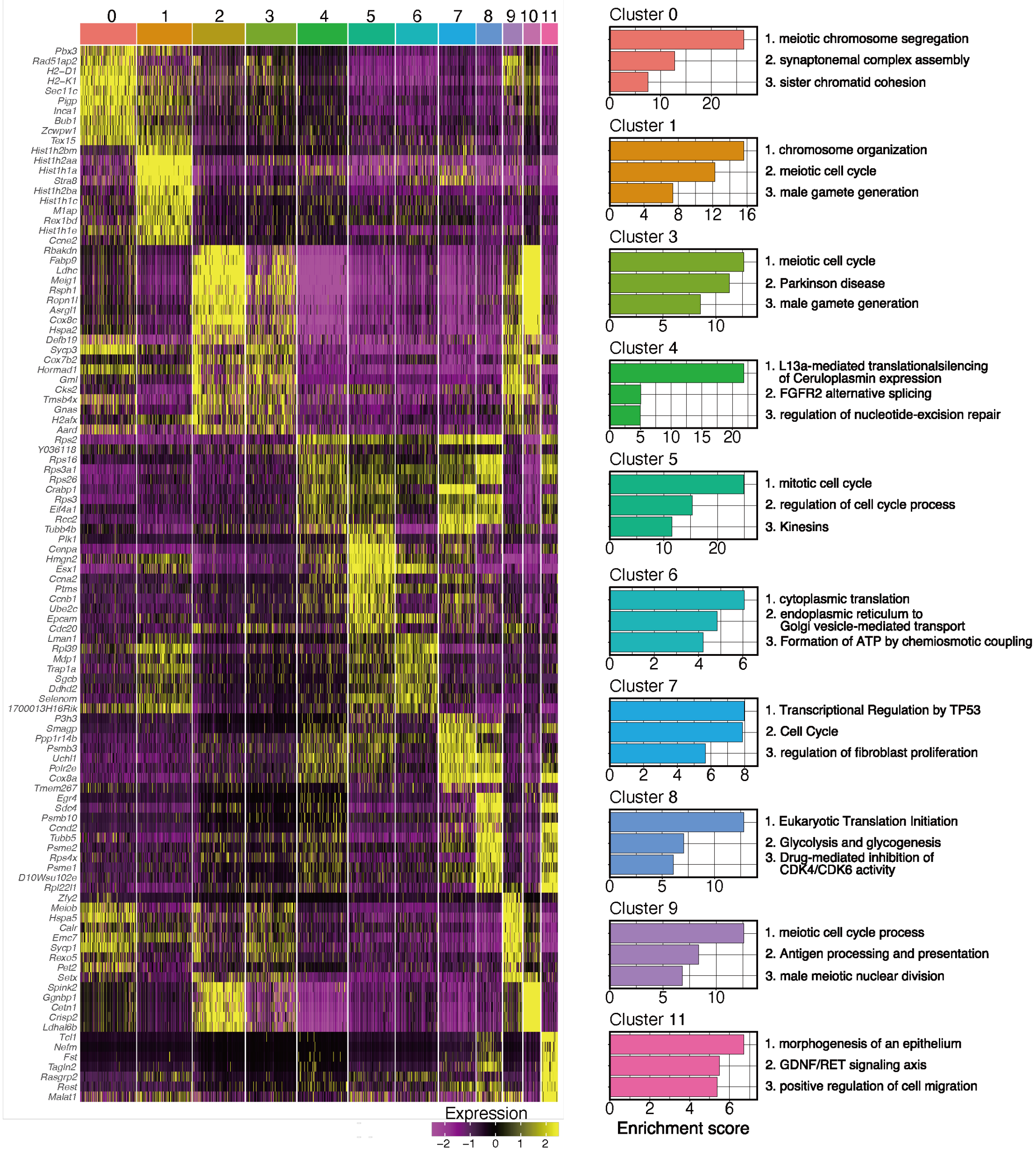


**Supplementary Figure 4. Clustering analysis of scRNA-seq data of WT and *Hsf5* KO spermatogenic germ cells (related to Figure 5)**

**(A)** Summary table of the 10X Genomics Chromium metrics for scRNA-seq analysis with WT and *Hsf5* KO testicular germ cells (P16). Indicated numbers of testes were pooled. Total number of testicular germ cells that were subjected to RNA-seq analysis and separated from other testicular somatic cells are shown. Percentages of the extracted testicular germ cells per total single cells that were subjected to RNA-seq analysis are shown.
**(B)** UMAP representation of scRNA-seq transcriptome profiles for testicular cells from P16 WT and *Hsf5* KO testes.
**(C)** Heat plot for the top 10 representative genes of each cluster. The relative expression levels are indicated with the color codes.
**(D)** Gene enrichment analysis of DEGs in the UMAP-defined cell clusters.

**A**

| Accession | Description | # AAs | HSF5-N1 IP (High salt) | | | HSF5-N2 IP (High salt) | | |
|---|---|---|---|---|---|---|---|---|
| | | | Score | PSMs | Area | Score | PSMs | Area |
| P17156 | Heat shock-related 70 kDa protein 2 [HSP72_MOUSE] | 633 | 573.65 | 34 | 4.729E7 | 1019.86 | 59 | 1.081E8 |
| Q9EPU4 | Cleavage and polyadenylation specificity factor subunit 1 [CPSF1_MOUSE] | 1441 | 25.54 | 1 | 1.067E6 | 705.52 | 49 | 5.726E7 |
| Q9EQK5 | Major vault protein [MVP_MOUSE] | 861 | 1265.42 | 72 | 5.394E7 | 622.00 | 48 | 2.272E7 |
| Q5ND04 | Heat shock factor protein 5 [HSF5_MOUSE] | 624 | 691.95 | 52 | 2.569E8 | 441.77 | 34 | 1.812E8 |
| P63017 | Heat shock cognate 71 kDa protein [HSP7C_MOUSE] | 646 | 218.90 | 17 | 3.756E7 | 434.93 | 24 | 9.073E7 |
| P68372 | Tubulin beta-4B chain [TBB4B_MOUSE] | 445 | 316.81 | 18 | 3.767E7 | 388.87 | 16 | 5.121E7 |
| Q03265 | ATP synthase subunit alpha, mitochondrial [ATPA_MOUSE] | 553 | 297.39 | 17 | 1.477E7 | 366.89 | 19 | 1.787E7 |
| P38647 | Stress-70 protein, mitochondrial [GRP75_MOUSE] | 679 | 312.83 | 12 | 1.249E7 | 333.10 | 16 | 1.745E7 |
| P99024 | Tubulin beta-5 chain [TBB5_MOUSE] | 444 | 271.78 | 14 | 3.290E7 | 318.63 | 14 | 4.575E7 |
| P20152 | Vimentin [VIME_MOUSE] | 466 | 137.88 | 12 | 1.698E7 | 304.21 | 18 | 1.879E7 |
| P60710 | Actin, cytoplasmic 1 [ACTB_MOUSE] | 375 | 295.67 | 17 | 4.417E7 | 302.22 | 17 | 5.940E7 |
| P05214 | Tubulin alpha-3 [TBA3_MOUSE] | 450 | 290.31 | 13 | 4.465E7 | 280.25 | 15 | 5.056E7 |
| P68369 | Tubulin alpha-1A [TBA1A_MOUSE] | 451 | 317.53 | 13 | 4.647E7 | 276.65 | 15 | 4.885E7 |
| Q9ERD7 | Tubulin beta-3 [TBB3_MOUSE] | 450 | 225.86 | 12 | 2.475E7 | 275.08 | 14 | 2.732E7 |
| P08113 | Endoplasmin [ENPL_MOUSE] | 802 | 179.22 | 10 | 6.059E6 | 243.51 | 9 | 7.597E6 |
| P56480 | ATP synthase subunit beta, mitochondrial [ATPB_MOUSE] | 529 | 308.82 | 16 | 2.095E7 | 240.35 | 16 | 2.534E7 |
| Q8K310 | Matrin-3 [MATR3_MOUSE] | 846 | 121.02 | 10 | 6.348E6 | 238.97 | 13 | 1.208E7 |
| Q8VEK3 | Heterogeneous nuclear ribonucleoprotein U [HNRPU_MOUSE] | 800 | 119.56 | 9 | 1.059E7 | 222.33 | 9 | 1.563E7 |
| Q91YQ5 | Dolichyl-diphosphooligosaccharide--protein glycosyltransferase subunit 1 [RPN1_MOUSE] | 608 | 144.81 | 7 | 7.731E6 | 216.28 | 10 | 1.286E7 |
| Q9D0E1 | Heterogeneous nuclear ribonucleoprotein M [HNRPM_MOUSE] | 729 | 137.64 | 10 | 9.755E6 | 186.96 | 9 | 1.766E7 |
| Q61584 | Fragile X mental retardation syndrome-related protein 1 [FXR1_MOUSE] | 677 | 139.85 | 9 | 1.097E7 | 127.79 | 6 | 6.214E6 |
| Q7TSG5 | SH3 domain-containing protein 21 [SH321_MOUSE] | 549 | 121.41 | 5 | 7.370E6 | 120.84 | 6 | 8.156E6 |
| Q8CGW4 | Transcription factor SOX-30 [SOX30_MOUSE] | 782 | 73.60 | 7 | 1.121E7 | 115.58 | 6 | 8.326E6 |
| Q9DA79 | Dipeptidase 3 [DPEP3_MOUSE] | 493 | 105.14 | 8 | 2.278E7 | 108.19 | 7 | 1.688E7 |
| O54724 | Polymerase I and transcript release factor [PTRF_MOUSE] | 392 | 191.19 | 8 | 1.823E7 | 104.32 | 3 | 6.814E6 |
| P11499 | Heat shock protein HSP 90-beta [HS90B_MOUSE] | 724 | 91.91 | 2 | 4.565E6 | 101.92 | 2 | 6.027E6 |
| P29341 | Polyadenylate-binding protein 1 [PABP1_MOUSE] | 636 | 51.03 | 1 | 4.204E6 | 98.73 | 3 | 3.601E6 |
| P70372 | ELAV-like protein 1 [ELAV1_MOUSE] | 326 | 65.26 | 2 | 4.341E6 | 96.54 | 2 | 6.476E6 |
| Q9JHU4 | Cytoplasmic dynein 1 heavy chain 1 [DYHC1_MOUSE] | 4644 | 62.00 | 5 | 2.529E6 | 91.14 | 8 | 3.770E6 |
| O70133 | ATP-dependent RNA helicase A [DHX9_MOUSE] | 1380 | 63.95 | 5 | 4.121E6 | 90.18 | 8 | 5.170E6 |
| P32037 | Solute carrier family 2, facilitated glucose transporter member 3 [GTR3_MOUSE] | 493 | 158.11 | 6 | 2.152E7 | 89.06 | 5 | 2.909E7 |
| O54734 | Dolichyl-diphosphooligosaccharide--protein glycosyltransferase 48 kDa subunit [OST48_MOUSE] | 441 | 61.91 | 2 | 3.362E6 | 82.39 | 3 | 5.284E6 |
| P62806 | Histone H4 [H4_MOUSE] | 103 | 45.36 | 5 | 5.000E6 | 78.86 | 5 | 4.503E6 |
| P27773 | Protein disulfide-isomerase A3 [PDIA3_MOUSE] | 505 | 33.29 | 2 | 3.625E6 | 78.22 | 4 | 5.319E6 |
| Q99PU5 | Long-chain-fatty-acid--CoA ligase ACSBG1 [ACBG1_MOUSE] | 721 | 29.75 | 2 | 2.054E6 | 78.03 | 3 | 2.064E6 |
| Q8CGW9 | Doublesex- and mab-3-related transcription factor C2 [DMRTD_MOUSE] | 370 | 70.36 | 5 | 9.144E6 | 75.37 | 3 | 3.111E6 |
| Q8BMS1 | Trifunctional enzyme subunit alpha, mitochondrial - [ECHA_MOUSE] | 763 | 30.93 | 2 | 2.175E6 | 74.97 | 3 | 2.057E6 |
| Q8C2Q3 | RNA-binding protein 14 [RBM14_MOUSE] | 669 | 26.55 | 1 | 2.314E6 | 74.90 | 4 | 5.640E6 |
| P14733 | Lamin-B1 [LMNB1_MOUSE] | 588 | 73.23 | 7 | 5.381E6 | 71.52 | 4 | 6.033E6 |
| P61407 | Tudor domain-containing protein 6 [TDRD6_MOUSE] | 2134 | 48.13 | 3 | 1.540E6 | 66.69 | 5 | 3.593E6 |
| Q8VIJ6 | Splicing factor, proline- and glutamine-rich [SFPQ_MOUSE] | 699 | 42.29 | 3 | 1.733E6 | 65.61 | 3 | 1.617E6 |
| P52194 | Calmegin [CLGN_MOUSE] | 611 | 48.34 | 2 | 4.603E6 | 65.50 | 3 | 7.966E6 |
| Q01320 | DNA topoisomerase 2-alpha [TOP2A_MOUSE] | 1528 | 126.00 | 7 | 7.386E6 | 64.92 | 2 | 5.669E6 |
| Q9DB77 | Cytochrome b-c1 complex subunit 2, mitochondrial [QCR2_MOUSE] | 453 | 33.38 | 1 | 2.741E6 | 62.94 | 2 | 7.272E6 |

| Accession | Description | # AAs | HSF5-N1 IP (High salt) | | | HSF5-N2 IP (High salt) | | |
|---|---|---|---|---|---|---|---|---|
| | | | Score | PSMs | Area | Score | PSMs | Area |
| P0CG49 | Polyubiquitin-B [UBB_MOUSE] | 305 | 55.68 | 4 | 2.408E7 | 61.72 | 5 | 2.220E7 |
| P57776 | Elongation factor 1-delta [EF1D_MOUSE] | 281 | 39.17 | 1 | 3.438E6 | 59.96 | 2 | 4.674E6 |
| P35564 | Calnexin [CALX_MOUSE] | 591 | 32.23 | 1 | 2.191E6 | 59.45 | 3 | 1.953E6 |
| O09106 | Histone deacetylase 1 [HDAC1_MOUSE] | 482 | 22.51 | 3 | 2.412E6 | 58.75 | 2 | 1.911E6 |
| Q61656 | Probable ATP-dependent RNA helicase DDX5 [DDX5_MOUSE] | 614 | 28.16 | 1 | 3.101E6 | 57.52 | 2 | 3.231E6 |
| Q3V132 | ADP/ATP translocase 4 [ADT4_MOUSE] | 320 | 40.21 | 2 | 3.137E6 | 56.24 | 2 | 3.792E6 |
| P10126 | Elongation factor 1-alpha 1 [EF1A1_MOUSE] | 462 | 60.69 | 3 | 1.866E7 | 52.64 | 2 | 1.279E7 |
| Q8VEM8 | Phosphate carrier protein, mitochondrial [MPCP_MOUSE] | 357 | 38.36 | 2 | 5.860E6 | 52.43 | 2 | 7.930E6 |
| O55143 | Sarcoplasmic/endoplasmic reticulum calcium ATPase 2 [AT2A2_MOUSE] | 1044 | 33.29 | 3 | 3.871E6 | 50.45 | 4 | 5.011E6 |
| Q60931 | Voltage-dependent anion-selective channel protein 3 [VDAC3_MOUSE] | 283 | 51.73 | 2 | 6.421E6 | 50.28 | 1 | 8.574E6 |
| O88569 | Heterogeneous nuclear ribonucleoproteins A2/B1 [ROA2_MOUSE] | 353 | 35.86 | 1 | 2.741E6 | 49.27 | 4 | 3.748E6 |
| Q0GGX2 | Zinc finger protein 541 [ZN541_MOUSE] | 1363 | 127.32 | 8 | 1.319E7 | 48.78 | 2 | 6.079E6 |
| Q60930 | Voltage-dependent anion-selective channel protein 2 [VDAC2_MOUSE] | 295 | 42.06 | 4 | 1.245E7 | 48.37 | 3 | 1.123E7 |
| Q6PDQ2 | Chromodomain-helicase-DNA-binding protein 4 [CHD4_MOUSE] | 1915 | 38.34 | 2 | 2.817E6 | 47.69 | 3 | 2.955E6 |
| Q9D1D4 | Transmembrane emp24 domain-containing protein 10 [TMEDA_MOUSE] | 219 | 34.42 | 1 | 0.000E0 | 47.15 | 3 | 3.238E6 |
| Q9JMB7 | Piwi-like protein 1 [PIWL1_MOUSE] | 862 | 31.88 | 1 | 8.271E5 | 46.26 | 4 | 2.922E6 |
| Q501J6 | Probable ATP-dependent RNA helicase DDX17 [DDX17_MOUSE] | 650 | 28.16 | 1 | 3.101E6 | 44.72 | 3 | 2.649E6 |
| Q921M3 | Splicing factor 3B subunit 3 [SF3B3_MOUSE] | 1217 | 28.20 | 1 | 1.762E6 | 44.40 | 2 | 2.322E6 |
| Q922R8 | Protein disulfide-isomerase A6 [PDIA6_MOUSE] | 440 | 34.98 | 1 | 1.568E6 | 42.26 | 2 | 2.322E6 |
| Q6P5H2 | Nestin[NEST_MOUSE] | 1864 | 48.87 | 4 | 4.569E8 | 41.67 | 3 | 1.485E8 |
| Q9Z2X1 | Heterogeneous nuclear ribonucleoprotein F [HNRPF_MOUSE] | 415 | 75.80 | 3 | 2.795E6 | 41.33 | 2 | 2.469E6 |
| P67778 | Prohibitin [PHB_MOUSE] | 272 | 34.98 | 2 | 1.200E6 | 40.50 | 2 | 1.477E6 |
| P00342 | L-lactate dehydrogenase C chain [LDHC_MOUSE] | 332 | 61.52 | 3 | 3.909E6 | 40.11 | 2 | 1.116E7 |
| Q9DBG6 | Dolichyl-diphosphooligosaccharide--protein glycosyltransferase subunit 2 [RPN2_MOUSE] | 631 | 25.71 | 1 | 4.215E6 | 39.28 | 2 | 7.582E6 |
| Q80X56 | E3 ubiquitin-protein ligase TRIM69 [TRI69_MOUSE] | 500 | 58.73 | 2 | 2.273E6 | 39.22 | 1 | 4.108E6 |
| E9Q7E2 | AT-rich interactive domain-containing protein 2 [ARID2_MOUSE] | 1828 | 30.10 | 2 | 3.943E6 | 39.08 | 5 | 4.280E6 |
| O35737 | Heterogeneous nuclear ribonucleoprotein H [HNRH1_MOUSE] | 449 | 103.28 | 3 | 3.429E6 | 38.38 | 2 | 3.179E6 |
| P01942 | Hemoglobin subunit alpha [HBA_MOUSE] | 142 | 27.35 | 2 | 4.142E6 | 36.53 | 1 | 4.191E6 |
| Q9Z1Q9 | Valine--tRNA ligase [SYVC_MOUSE] | 1263 | 61.71 | 4 | 1.597E6 | 36.46 | 2 | 1.771E6 |
| E9PYK3 | Protein mono-ADP-ribosyltransferase PARP4 [PARP4_MOUSE] | 1969 | 131.03 | 9 | 9.979E6 | 36.17 | 1 | 2.839E6 |
| P61979 | Heterogeneous nuclear ribonucleoprotein K [HNRPK_MOUSE] | 463 | 26.48 | 1 | 1.618E6 | 35.93 | 2 | 3.170E6 |
| Q562E2 | BTB/POZ domain-containing protein KCTD19 [KCD19_MOUSE] | 927 | 72.33 | 8 | 4.212E6 | 34.09 | 2 | 6.260E6 |
| Q9CZ13 | Cytochrome b-c1 complex subunit 1, mitochondrial [QCR1_MOUSE] | 480 | 34.85 | 2 | 5.214E6 | 34.07 | 2 | 5.291E6 |
| A2A8L1 | Chromodomain-helicase-DNA-binding protein 5 [CHD5_MOUSE] | 1946 | 38.34 | 2 | 2.817E6 | 33.63 | 2 | 4.662E6 |
| Q8BFZ9 | Erlin-2 [ERLN2_MOUSE] | 340 | 37.09 | 2 | 2.016E6 | 33.60 | 3 | 2.265E6 |
| Q91Z31 | Polypyrimidine tract-binding protein 2 [PTBP2_MOUSE] | 531 | 24.81 | 2 | 3.801E6 | 32.71 | 1 | 2.418E6 |
| P01029 | Complement C4-B [CO4B_MOUSE] | 1738 | 21.50 | 2 | 4.109E6 | 31.08 | 1 | 9.342E6 |
| Q99KI0 | Aconitate hydratase, mitochondrial [ACON_MOUSE] | 780 | 43.86 | 2 | 2.998E6 | 30.51 | 1 | 4.221E6 |
| Q8BMF4 | Dihydrolipoyllysine-residue acetyltransferase component of pyruvate dehydrogenase complex, mitochondrial | 642 | 43.70 | 4 | 5.157E6 | 29.63 | 2 | 2.550E6 |
| P10853 | Histone H2B type 1-F/J/L [H2B1F_MOUSE] | 126 | 35.81 | 2 | 2.211E6 | 29.44 | 1 | 6.213E6 |
| Q91ZW3 | SWI/SNF-related matrix-associated actin-dependent regulator of chromatin subfamily A member 5 | 1051 | 41.72 | 3 | 3.472E6 | 28.24 | 1 | 5.648E6 |
| P97496 | SWI/SNF complex subunit SMARCC1 [SMRC1_MOUSE] | 1104 | 31.93 | 2 | 1.165E6 | 22.46 | 2 | 1.536E6 |
| P17225 | Polypyrimidine tract-binding protein 1 [PTBP1_MOUSE] | 555 | 31.94 | 2 | 2.602E6 | 14.81 | 2 | 2.813E6 |

**B**

| Accession | Description | # AAs | HSF5-C IP (High salt) | | |
|---|---|---|---|---|---|
| | | | Score | PSMs | Area |
| Q5ND04 | Heat shock factor protein 5 [HSF5_MOUSE] | 624 | 1772.34 | 122 | 7.499E8 |
| Q8K310 | Matrin-3 OS=Mus musculus [MATR3_MOUSE] | 846 | 340.79 | 19 | 3.785E7 |
| P38647 | Stress-70 protein [GRP75_MOUSE] | 679 | 161.58 | 6 | 1.091E7 |
| Q501J6 | Probable ATP-dependent RNA helicase DDX17 [DDX17_MOUSE] | 650 | 145.70 | 6 | 9.757E6 |
| Q8VEK3 | Heterogeneous nuclear ribonucleoprotein U [HNRPU_MOUSE] | 800 | 110.11 | 4 | 2.389E7 |
| Q9WTM5 | RuvB-like 2 [RUVB2_MOUSE] | 463 | 95.34 | 3 | 1.161E7 |
| Q01320 | DNA topoisomerase 2-alpha [TOP2A_MOUSE] | 1528 | 88.84 | 5 | 1.503E7 |
| Q8C2Q3 | RNA-binding protein 14 [RBM14_MOUSE] | 669 | 71.11 | 3 | 1.125E7 |
| P29341 | Polyadenylate-binding protein 1 [PABP1_MOUSE] | 636 | 58.12 | 2 | 1.526E7 |
| Q8BL97 | Serine/arginine-rich splicing factor 7 [SRSF7_MOUSE] | 267 | 55.37 | 2 | 1.317E7 |
| O54941 | SWI/SNF-related matrix-associated actin-dependent regulator of chromatin subfamily E member 1 | 411 | 54.56 | 2 | 6.770E6 |
| P62806 | Histone H4 [H4_MOUSE] | 103 | 52.87 | 4 | 8.472E6 |
| Q3TWW8 | Serine/arginine-rich splicing factor 6 [SRSF6_MOUSE] | 339 | 46.24 | 2 | 3.030E6 |
| Q91ZW3 | SWI/SNF-related matrix-associated actin-dependent regulator of chromatin subfamily A member 5 | 1051 | 44.64 | 4 | 7.137E6 |
| Q6PDG5 | SWI/SNF complex subunit SMARCC2 [SMRC2_MOUSE] | 1213 | 41.73 | 4 | 9.360E6 |
| Q9Z2X1 | Heterogeneous nuclear ribonucleoprotein F [HNRPF_MOUSE] | 415 | 39.05 | 2 | 4.781E6 |
| P14733 | Lamin-B1 [LMNB1_MOUSE] | 588 | 36.78 | 2 | 1.347E7 |
| Q9Z204 | Heterogeneous nuclear ribonucleoproteins C1/C2 [HNRPC_MOUSE] | 313 | 34.46 | 2 | 3.880E6 |
| Q562E2 | BTB/POZ domain-containing protein KCTD19 [KCD19_MOUSE] | 927 | 30.34 | 2 | 4.025E6 |
| Q60749 | KH domain-containing, RNA-binding, signal transduction-associated protein 1 [KHDR1_MOUSE] | 443 | 24.09 | 3 | 5.713E6 |

**Supplementary Figure 5. MS analyses of HSF5 interacting factors in testis extracts (related to Figure 6)**

**(A)**The immunoprecipitates (IP) by two different HSF5-N antibodies (HSF5-N1, HSF5-N2) from the chromatin-bound fractions of the testis extracts were subjected to liquid chromatography tandem-mass spectrometry (LC-MS/MS) analyses. The proteins

identified by the LC-MS/MS analysis of HSF5N-IP are presented after excluding the proteins detected in the control IgG-IP. The proteins that were detected in both HSF5-N1 IP and HSF5-N2 IP with more than 1 different peptide hits are listed with SwissProt accession number, Amino acid length, the number of peptide hits and Mascot scores.
**(B)**The immunoprecipitates (IP) by HSF5-C antibody from the chromatin-bound fractions of the testis extracts. The proteins identified by the LC-MS/MS analysis of HSF5C-IP are presented as in A. See also Table S5 for the raw data of LC-MSMS analysis.
Note that SMARCA5 and KCTD19 was repeatedly identified in the immunoprecipitates by HSF5-N1, HSF5-N2 HSF5-C antibodies.

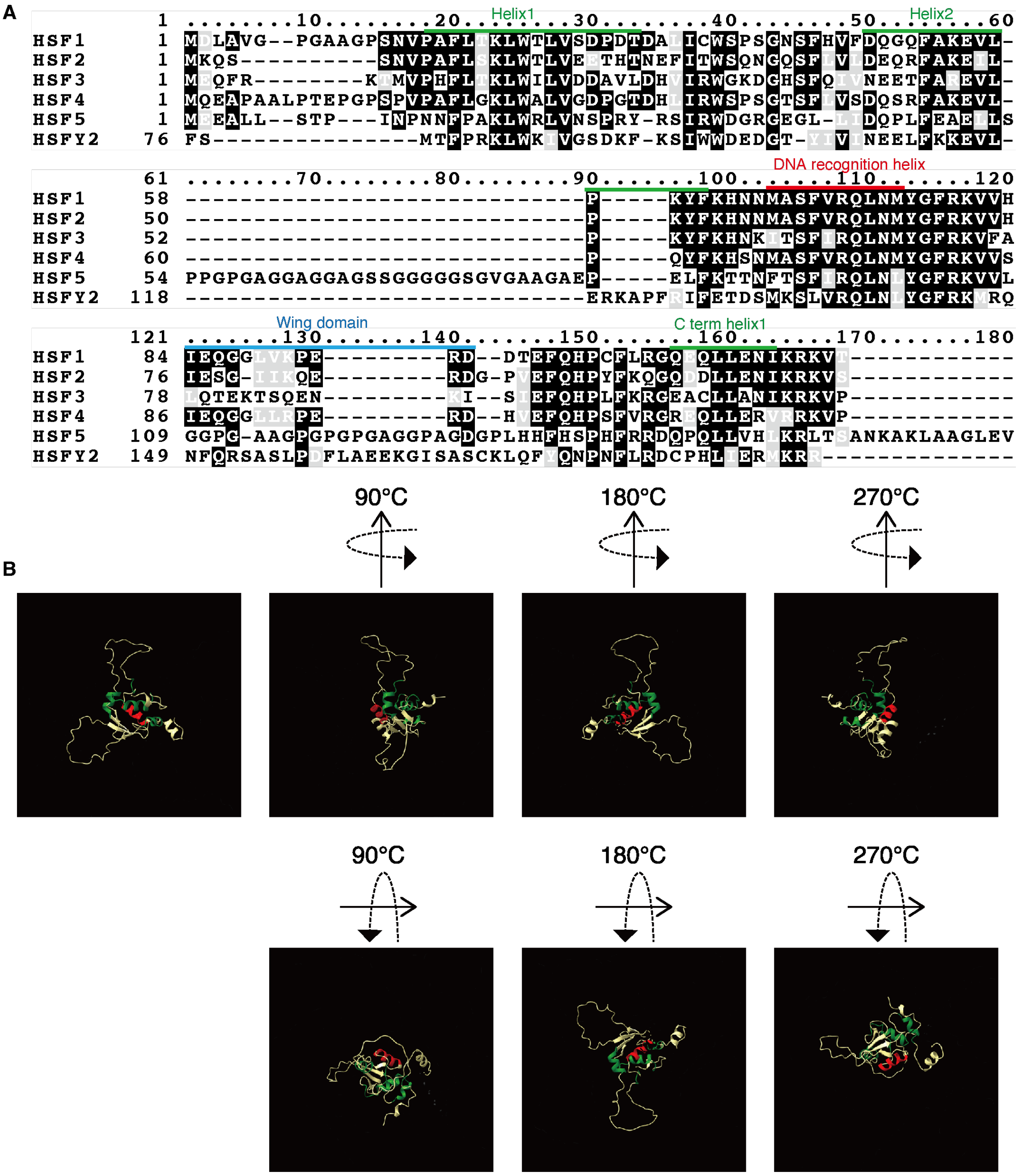


**Supplementary Figure 6. Sequence alignment of DNA-binding domain of the mouse HSFs (related to Figure 6)**

**(A)** Amino acid sequences of DNA-binding domain in the mouse HSFs are aligned. HSF1, HSF2, and HSF4 possess a DNA recognition helix (red), containing a conserved Ser-Phe-Val-Arg-Gln amino acid sequence, which is known to insert into the major groove of the HSE. HSF5 possesses a Ser-Phe-Ile-Arg-Gln amino acid sequence at the corresponding position. HSF5 possesses insertion of amino acid sequence between the helix 2 and DNA recognition helix.

**(B)** Ribbon models of HSF5-DBD (1-167 a.a.) is shown that was predicted from AlphaFold2. Helixes are colored as shown in (A).

Figure 1D

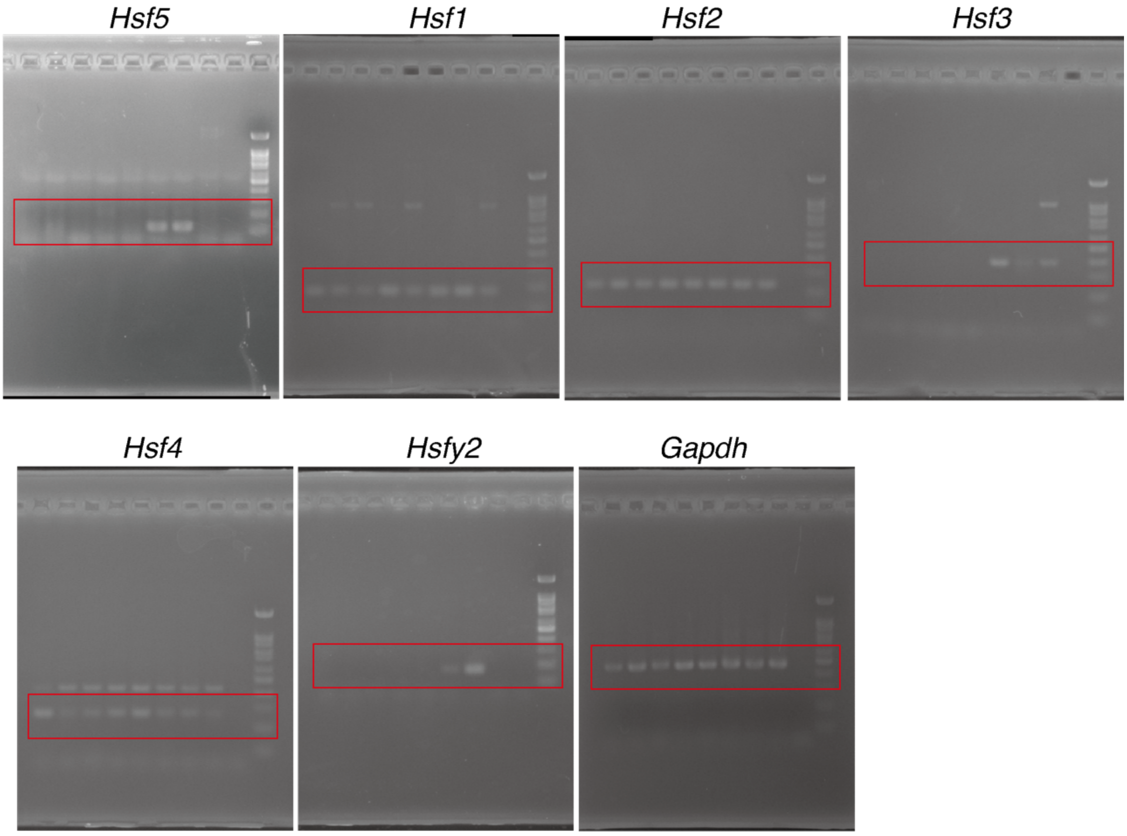


Figure 2B

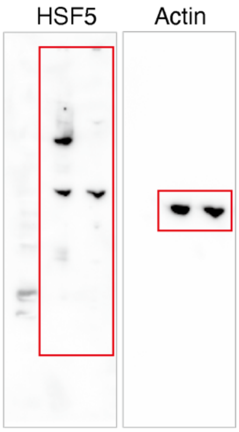


Figure 6G

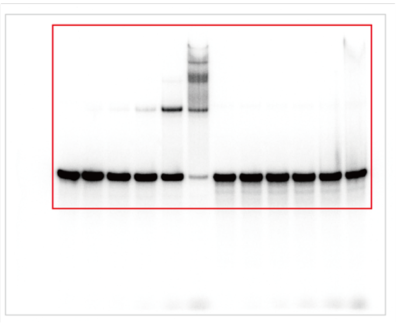

Figure 6H

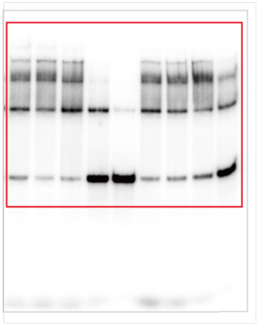

**Supplementary Figure 7. Uncropped images of gels and blots. (related to Figure 1, 2, 6)**

Full-length / uncropped images of agarose gel (Fig1D), western blot (Fig2B) and the scanned images of autoradiograph (Fig6G, Fig6H) are shown.
